# High-throughput cultivation of stable, diverse, fecal-derived microbial communities to model the intestinal microbiota

**DOI:** 10.1101/2020.07.06.190181

**Authors:** Andrés Aranda-Díaz, Katharine Michelle Ng, Tani Thomsen, Imperio Real-Ramírez, Dylan Dahan, Susannah Dittmar, Carlos Gutierrez Gonzalez, Taylor Chavez, Kimberly S. Vasquez, Taylor H. Nguyen, Feiqiao Brian Yu, Steven K. Higginbottom, Norma F. Neff, Joshua E. Elias, Justin L. Sonnenburg, Kerwyn Casey Huang

## Abstract

Mechanistic understanding of the impacts of the gut microbiota on human health has been hampered by limited throughput in animal models. To enable systematic interrogation of gut-relevant microbial communities, here we generated hundreds of *in vitro* communities cultured from diverse stool samples in various media. Species composition revealed stool-derived communities that are phylogenetically complex, diverse, stable, and highly reproducible. Community membership depended on both medium and initial inoculum, with certain media preserving inoculum compositions. Different inocula yielded different community compositions, indicating their potential for personalized therapeutics. Communities were robust to freezing and large-volume culturing, enabling future translational applications. Defined communities were generated from isolates and reconstituted growth and composition similar to those of communities derived from stool inocula. Finally, *in vitro* experiments probing the response to ciprofloxacin successfully predicted many changes observed *in vivo*, including the resilience and sensitivity of each *Bacteroides* species. Thus, stool-derived *in vitro* communities constitute a powerful resource for microbiota research.

## Introduction

The gut microbiota is a diverse community that performs functions important for host physiology, including interactions with the immune system (Hooper et al., 2012), metabolism (Sonnenburg and Backhed, 2016), and the nervous system (Mayer et al., 2014). A wide range of environmental perturbations, including diet (Kashyap et al., 2013), antibiotics (Schubert et al., 2015), and osmotic diarrhea (Tropini et al., 2018), lead to changes in community composition and function. Human and animal models have enabled some mechanistic understanding of these responses. For example, gnotobiotic mouse models provide insights into particular microbial communities (Becker et al., 2011; Faith et al., 2011; Mark Welch et al., 2017; Reyes et al., 2013; Rezzonico et al., 2011; Turnbaugh et al., 2009) composed of a single species, a defined set of species, or of complex samples originating from specific hosts such as humans.

Importantly, our understanding of microbiota recovery from antibiotics is limited. Antibiotics disrupt colonization resistance and lead to pathogen expansion and fecal shedding (Barthel et al., 2003; Doorduyn et al., 2006; Lawley et al., 2008; Pavia et al., 1990; Stecher et al., 2007), and can affect host physiology, including changes in adiposity, insulin resistance, and cognitive function (Cho et al., 2012; Cox et al., 2014; Frohlich et al., 2016; Hwang et al., 2015). Understanding antibiotic effects across microbiotas, doses, and treatment regimens will assist in mitigating these adverse effects. We recently showed, using mice colonized with human feces, that the microbiota shifts into a new, lower-diversity steady state after ciprofloxacin treatment, although certain taxa displayed a high level of resilience during recovery (Ng et al., 2019). It is unclear to what extent the response of each taxon can be predicted from its behavior in isolation; microbial interactions and drug modification can impact the efficacy of antibiotic treatment (Adamowicz et al., 2018; de Vos et al., 2017; Nicoloff and Andersson, 2016; Sanchez-Vizuete et al., 2015), which may underlie our current inability to explain individualized responses to fluoroquinolone treatment in humans (Dethlefsen and Relman, 2011) or to understand why some antibiotics, like vancomycin, negatively impact taxa known to be resistant to the drug *in vitro* (Ivanov et al., 2008).

*In vivo* investigations of the gut microbiota of humans and mice are limited in throughput, and it is often difficult to isolate and quantitatively tune the magnitude of perturbations. As a complementary approach, synthetic *in vitro* co-culturing of a defined community of isolated species has revealed key interspecies interactions and metabolic roles (de Vos et al., 2017; Gutierrez and Garrido, 2019; Kehe et al., 2019; Venturelli et al., 2018). However, such bottom-up approaches generally lack the complexity of typical gut microbiotas; further, microbes coexisting in a host shape each other’s evolutionary paths (Barroso-Batista et al., 2020), which could underlie individualized responses of the human gut microbiota to antibiotics (Dethlefsen and Relman, 2011). It is difficult to select a particular set of species to model complex ecosystems such as the mammalian gut without *a priori* knowledge of interspecies interactions; most approaches therefore choose representative species from the major taxa.

While sophisticated continuous-flow culture models (Macfarlane et al., 1998; Minekus et al., 1999; Van de Wiele et al., 2015) permit long time-scale experimentation and the manipulation of key environmental variables, these systems are also limited in throughput (to date, mini-bioreactor arrays have included 48 reactors running in parallel; (Auchtung et al., 2016). Other approaches to study the function of complex microbiotas *in vitro* include *ex vivo* resuspensions of stool or other natural communities in liquid media, for example to analyze drug metabolism by highly complex communities (Zimmermann et al., 2019). These approaches are higher throughput, but the extent to which they recapitulate the gut environment is not clear, and they have yet to be used to systematically study community assembly and growth from gut samples. In a recent study, diverse microbiomes from leaf and soil samples were passaged in minimal media with a single carbon source (Goldford et al., 2018). Batch culturing led to the selection of simple communities whose compositions were predictably governed by nutrient availability at the family but not the species level, revealing alternative species-level community assembly patterns (Goldford et al., 2018). The extent to which these findings extend to human-relevant microbiotas and more complex media is unknown.

To close these gaps, here we established that top-down, stool-derived *in vitro* communities (SICs) are powerful tools for modeling responses of the gut microbiota to perturbations *in vivo*. We demonstrated that hundreds of SICs generated from distinct initial inocula in diverse media can be phylogenetically complex and diverse, and preserve the structure of the initial inoculum with generally stable and reproducible compositions. SICs can be frozen, grown in larger volumes, and reconstituted from isolates without affecting composition. Most importantly, upon antibiotic treatment, many of the responses of the SICs mimicked changes observed in the antibiotic-treated humanized gnotobiotic mice from which the initial inocula were sampled, indicating that SICs could be used to predict and interpret the results of *in vivo* experiments in high throughput.

## Results

### Diet and antibiotic treatment provide a diverse pool of inocula

To generate SICs in a straightforward, scalable, and reproducible manner, we focused on ex-germ-free mice colonized with human feces. These “humanized” mice facilitate controlled experimental manipulation within a mammalian host, enabling comparisons between *in vitro* and *in vivo* behaviors of communities of human commensals. We inoculated humanized mouse fecal samples into various media and repeatedly passaged in anaerobic batch cultures (Methods). We sought to inoculate with a set of fecal samples that varied widely in composition, yet were composed of the same set of members so that they could be readily compared.

We took advantage of our previous findings that diet and antibiotics have large, distinct, interacting effects on the composition of the microbiota (Ng et al., 2013; Sonnenburg et al., 2016). We inoculated 20 germ-free mice (5 per cage) with feces from a single human donor and allowed the microbiota to equilibrate for 6 weeks to allow for mucus normalization (Johansson et al., 2015). We shifted half of the mice (2 cages) from a standard diet (SD) to one deficient in microbiota-accessible carbohydrates (MD) and allowed the microbiota to re-equilibrate for two weeks. We then gavaged all mice (2 cages SD, 2 cages MD; *n*=20 total) for 5 days twice daily with 3 mg ciprofloxacin, which inhibits bacterial type II topoisomerases such as *Escherichia coli* DNA gyrase. Since we previously showed that the responses of cage-mates to antibiotics are highly reproducible and stereotypical (Ng et al., 2019), we collected fecal samples from two randomly selected mice in different cages from each dietary condition on day 0 (pre-treatment) and on day 1 (peak of treatment, when the culturable load reached a maximum decrease of 10- to 100-fold (Ng et al., 2019)), day 5 (residual treatment, when ciprofloxacin was not being administered but was still detected in feces (Ng et al., 2019)), and day 14 (post-treatment) (Fig. 1A).

**Figure 1:**
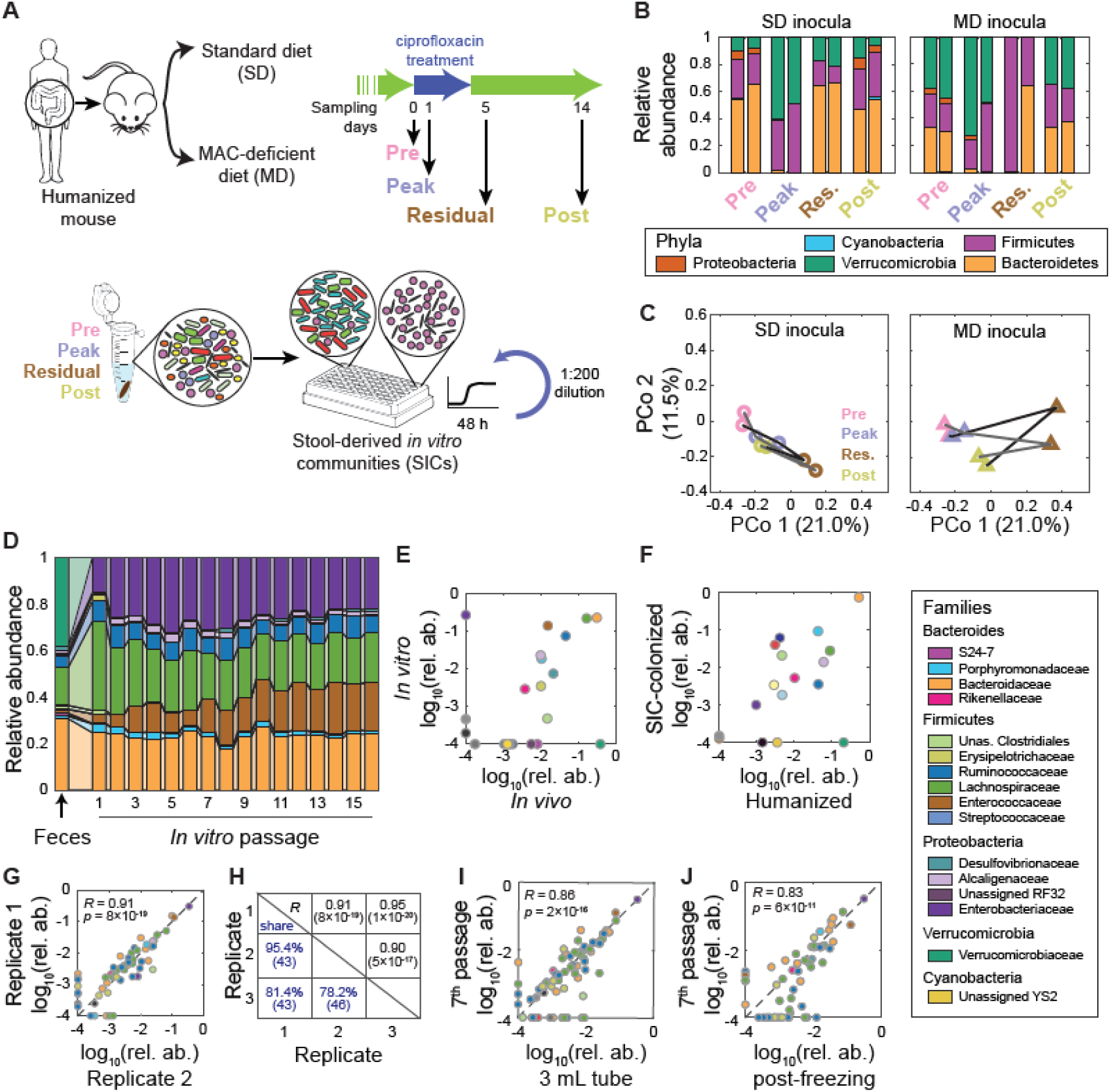
High-throughput cultivation of feces from humanized mice yields stable, complex and reproducible microbial communities. A) Experimental setup. Germ-free mice colonized with feces from a single human donor (“humanized”) were fed a SD or an MD and treated with ciprofloxacin for 5 days. Inocula from two mice on each diet were taken on four days (0, 1, 5, 14) before, during, and after treatment, and were inoculated into anaerobic batch culture and passaged with dilution every 48 h to derive SICs. Sixteen samples (2 diets, 2 mice, 4 time points during ciprofloxacin treatment) were inoculated into 4 media (BHI, TYG, GAM, and YCFA) in triplicate. B) Ciprofloxacin elicits large changes in microbiota composition at the phylum level *in vivo*. Each time point has two bars corresponding to the two mice in each group. C) Diet alters the trajectory of microbiota reorganization during ciprofloxacin treatment *in vivo*. PCoA of community composition of the fecal inocula in (B) using unweighted Unifrac distance computed on all *in vivo* and *in vitro* samples at the ASV level. D) *In vitro* passaging produces stable and complex SICs. Family-level compositions of an example SIC derived from the fecal inoculum of pre-treatment MD mouse 1 during 16 rounds of *in vitro* passaging in BHI. E) *In vitro* passaging can produce an SIC that resembles the fecal inoculum. Mean family-level log_10_(relative abundance) of passages 4-16 for the SIC in (D) and the fecal inoculum from which it originated. Families with relative abundance of 0 were set to 10^-4^ for visualization. F) Colonization of germ-free mice with an SIC restores families overrepresented *in vitro* back to levels similar to those in the humanized mice from which the SIC was derived. Comparison of the family-level log_10_(relative abundance) of a humanized-mouse fecal inoculum and the mean family-level log_10_(relative abundance) of ex-germ-free mice colonized with an SIC derived from the humanized-mouse fecal inoculum (inoculum *n*=1, SIC-colonized *n*=3). Families with relative abundance of 0 were set to 10^-4^ for visualization. G,H) *In vitro*-passaged SICs are highly reproducible. (G) ASV-level log_10_(relative abundance) for two technical replicates of the SIC in (D) after 7 passages. ASVs with relative abundance of 0 were set to 10^-4^ for visualization. (H) Summary of the reproducibility of the three replicates for the SIC in (D). *R* and *p* were computed using only ASVs present in both samples. Also shown is the percentage of ASVs that are present in both replicates at >0.1% abundance (“shared”), with the total number of ASVs in parenthesis. I,J) SICs can be grown in larger volumes and can be frozen and revived without affecting composition. ASV-level log_10_(relative abundance) values for the SIC in (D) after 7 passages were compared with the same SIC grown for one passage in a larger vessel without shaking (I), and after freezing with 25% glycerol at −80 °C and reviving for one passage (J). *R* and *p* were computed using only ASVs present in both samples. ASVs with relative abundance of zero were set to 10^-4^ for visualization.

Using 16S sequencing and analysis (Methods), we quantified the relative abundances of amplicon sequence variants (ASVs, or unique sequences) and mapped them to taxa. SD mice underwent a substantial change in fecal-sample composition due to ciprofloxacin treatment, with consistent reductions in Bacteroidetes and Proteobacteria and expansions in Firmicutes and Verrucomicrobia at the peak of treatment (Fig. 1B, S1A). The relative abundance of the Bacteroidetes mostly returned to pre-treatment levels by day 5, while the Proteobacteria and Verrucomicrobia did not fully recover until day 14 (Fig. 1B, S1A). Fecal samples from MD mice showed similar trends but had lower levels of Bacteroidetes throughout the experiment (Fig. 1B, S1A).

For SD mice, the number of uniquely detected ASVs (richness) dropped from 91 to 68 on day 1 of treatment and to 19 by day 5 (Fig. S1B). Post-treatment diversity partially recovered to 56 ASVs (Fig. S1B). Richness dynamics were qualitatively similar for MD mice (Fig. S1B).

Principal coordinates analysis (PCoA, Methods) indicated that microbiota trajectories were distinct between dietary conditions (Fig. 1C). Thus, feces from humanized mice fed distinct diets and sampled at different timepoints during antibiotics treatment yielded an array of qualitatively different inocula. All these inocula were derived from the same human donor sample, enabling the generation of SICs with distinct starting compositions but with the potential for overlapping ASVs, facilitating the assessment of culturing conditions on a common pool of microbes.

### *In vitro* cultivation of stable SICs from fecal samples

To generate SICs, we sampled mouse feces and quickly (<30 min after fecal pellet production) resuspended these 16 samples (2 diets, 4 treatment days, 2 mice per diet type) into media in anaerobic conditions and grew them in microplates in a plate reader at 37 °C (Methods, Fig. 1A). To explore multiple nutrient environments, we selected four common growth media: Brain Heart Infusion (BHI), Tryptone-Yeast extract-Glucose (TYG), Gifu Anaerobic Medium (GAM), and Yeast extract-Casitone-Fatty acids (YCFA). We inoculated three technical replicates of each resuspended fecal sample into wells of each medium to test the replicability of the resulting SICs.

Nutrient supplies differ over time in the gut; hence, we grew SICs in anaerobic batch culture rather than in a chemostat-like environment. Since some commensals require >24 h to reach saturation (Tramontano et al., 2018), we allowed all cultures to grow for 48 h, before diluting them 200-fold into fresh medium. We repeated this cycle for 2-4 weeks (7-14 passages), and after each sub-culturing, we split the samples; half were dedicated to glycerol frozen stocks and half to DNA extraction for 16S sequencing. Passaged SICs were not frozen or exposed to oxygen at any point, and aside from the brief period required for dilution, all samples were maintained at 37 °C throughout passaging.

To determine whether SICs reached stable compositions, we first focused on 16S data for the six SICs derived from pre-treatment samples from MD mice (Pre-MD) that were grown in BHI. These samples underwent 16 passages rather than the typical 7, which enabled us to identify whether changes in composition would occur over longer timescales. At the family level, SICs stabilized by the second passage (Fig. 1D) and retained most families from the original fecal sample (Fig. 1E). Of the families that were present in the inoculum but not in the passaged SIC, most were at low abundance *in vivo* (Fig. 1E); the lone exception was the Verrucomicrobiaceae, which were replaced by Enterobacteriaceae in the SIC (Fig. 1D,E).

To quantify the relevance of SICs as a model for the gut microbiota, we introduced an SIC into germ-free mice. Families that were overrepresented *in vitro* receded *in vivo*, with family-level abundances similar to those of the humanized mice from which the SIC was derived (Fig. 1F, Fig. S1C). To evaluate effects on host physiology we quantified host and microbial proteins in the feces of three groups of mice: humanized mice, germ-free mice, and SIC-colonized mice. The host proteomes of SIC-colonized mice were more similar to those of humanized mice than to those of germ-free mice, and immunoglobulins were upregulated after colonization in SIC-colonized mice (Fig. S1D). In the SIC metaproteome, proteins from two of the most abundant bacterial species (determined via 16S sequencing) were present at levels similar to those in SIC-colonized and humanized mice (Fig. S1E). These findings demonstrate that top-down *in vitro* cultivation from fecal samples can deterministically select complex SICs that are stable under passaging for several weeks; re-introduction of SICs into a host re-establishes gut microbiota composition and promotes host homeostasis.

A previous soil and plant microbiota passaging study reported that despite convergence at the family level, genus-level composition was highly divergent across technical replicates (Goldford et al., 2018). In our experiments, family-level dynamics were similar across all three replicate SICs from each of the two mice (Fig. S1F), as confirmed via PCoA (Fig. S1G). To test whether family-level replicability extended to the ASV level, we calculated the pairwise correlation coefficients of log_10_(relative abundance) between 7th passages of the 3 technical replicates of the SIC in Fig. 1D. Correlation coefficients and the percentage of ASVs shared (above 0.1% in both replicates) were uniformly high (Fig. 1G,H), and all ASVs that were present in only one replicate had relative abundance <1%. The three replicates contained 50 ASVs in total at >0.1%, which we define as a conservative measure of detectability, of which 14% were shared by exactly two replicates and 70% were shared by all three replicates. The 35 shared ASVs belonged to 9 families, 6 classes, and 3 phyla, and accounted for 97.5±1.1% (mean±standard deviation) of SIC total abundance. Hence, SIC replicability extends even to individual ASVs.

SICs were also reproducible across media and inocula (Fig. S1H, Supplementary Text). The lowest reproducibility was within SICs cultured in GAM and YCFA (Fig. S1I), where the dominance or absence of the Enterococcaceae explained most of the variation (Fig. S1J, Supplementary Text). Complex synthetic media mainly composed of amino acids and simple carbohydrates also drove the composition of SICs to be dominated by Enterococcaceae or Enterobacteriaceae (Fig. S1K). These findings indicate that *in vitro* culturing can lead to the selection of a few species in some media, and that SICs are sometimes stochastic, even in media such as YCFA that promote the growth of many commensals in isolation. These data also show that the SICs are generally diverse and reproducible in BHI and TYG.

For steady-state SICs derived from feces to be useful for further experimentation, they would ideally maintain their composition after freezing, and during growth in larger volumes and without shaking. We inoculated the frozen stock from the 7^th^-passage SIC in Fig. 1D (the last passage for which all other SICs in this work had frozen stocks, and a passage in which all BHI communities had achieved stability, Fig. S1L) and grew it in a 15-fold larger volume (3 mL) of BHI in glass test tubes without shaking. The tube-grown SIC had similar composition as the shaken SIC grown in a 96-well plate (Fig. 1I). Thus, shaking is not required, allowing a large number of plates to be grown in an incubator simultaneously, and large cultures can be readily produced.

To test SIC robustness to freezing, we inoculated −80 °C frozen stocks of the 3 mL test-tube cultures directly into fresh BHI and passaged them for three additional 48-h cycles. A single passage was sufficient to re-establish a composition closely matching that of the 7th passage pre-freezing (Fig. 1J); 37 of 50 ASVs at >0.1% were retained and accounted for 94.6% of the relative abundance pre-freezing. The composition was unchanged after two additional passages (Fig. S2). All other tested communities grown in BHI from other fecal inocula behaved similarly (Fig. S2), indicating that most taxa are preserved during freezing and thawing and that there were no substantive differential effects of freezing across taxa. Hence, SICs can be grown across a range of volumes, frozen, and revived without impact to the composition.

### Initial inoculum composition and nutrient conditions affect SIC composition

To determine how quickly SICs converge, we computed the weighted Unifrac distance in ASV abundances (Methods) and determined when the distance decreased below 0.15 (the mean weighted Unifrac distance between technical replicates at the third passage). While a few SICs with low replicability also exhibited longer convergence timescales, the vast majority converged to a steady state within 4 passages (Fig. S1L).

To determine the extent to which the abiotic environment and inoculum dictated steady-state SIC compositions after 7 passages, we performed PCoA of our 16S data for all passages of all SICs in all media. The first coordinate was largely determined by the presence or absence of several co-occurring families, including Enterobacteriaceae (Fig. S3A). Most of these families were present in the pre-treatment inocula and were undetectable in residual treatment (day 5) inocula (Fig. S3B), suggesting that most of the variation is explained by the fecal inocula, not the culture media. Similar to previous studies of communities inoculated from soil and leaf samples (Goldford et al., 2018), our data approximately clustered in the space of the first two principal coordinates according to culture media; BHI and TYG SICs overlapped but were well separated from GAM and even more so from YCFA (Fig. 2A). BHI, TYG, and GAM SICs were more similar to their inocula than YCFA SICs when using a weighted approach that accounts for taxonomical relative abundances (Fig. 2B). Using presence and absence of taxa as a measure of similarity (thus giving more weight to low-abundance taxa), BHI and TYG are better mimics of the conditions that produced the fecal pellet (Fig. 2A), regardless of the inoculating composition. The data also clearly clustered according to the time during antibiotic treatment when the inocula were sampled (Fig. 2A). Some SICs derived from the peak of antibiotic disturbance exhibited the largest differences from starting inoculum (Fig. 2A,C); instead, they more closely resembled SICs derived from residual treatment samples (Fig. 2A), likely due to low viable counts from samples at the peak of ciprofloxacin disturbance (Ng et al., 2019), implying that after antibiotic treatment, a fecal sample contains many non-viable cells. The direction separating clusters of sampling time was approximately orthogonal to the direction separating SICs by medium (Fig. 2A), indicating that both the availability of taxa in the inoculum and the ability of those taxa to grow as communities in a given medium dictate the composition of SICs.

**Figure 2:**
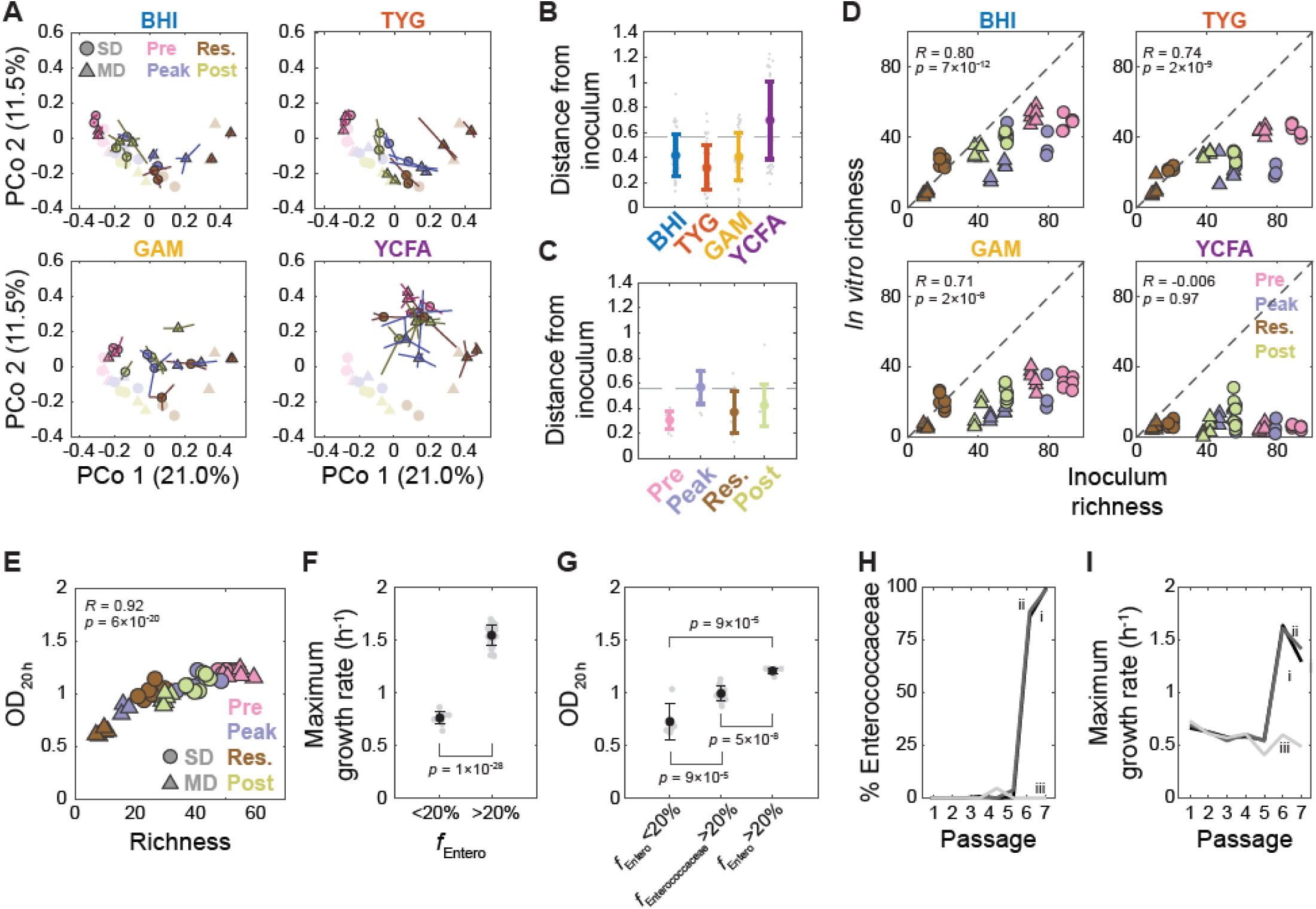
SIC composition is determined by inoculum and media, and is predictive of yield and growth rate. A) Medium and inoculum determine the final composition of passaged SICs. The 7^th^ passage of all 192 SICs in a PCoA of SIC composition using unweighted Unifrac distance computed on all *in vivo* and *in vitro* samples at the ASV level. Samples are separated by media, with colors and shapes representing the timepoint during ciprofloxacin treatment and diet, respectively, in the mice from which the inocula were taken. Symbols are the centroid of three replicates, with lines connecting the replicates to the centroid. Original fecal inocula are plotted in light colors (same data as in Fig. 1C). B) Most steady-state SICs are similar to the fecal samples from which they were derived. Weighted Unifrac distance of the 7^th^ passage of each SIC to the corresponding fecal inoculum. Colored circles, mean distance for each medium; individual SICs in gray. Error bars, standard deviations (SD); *n*=48. Dashed line, mean distance between fecal samples. C) SICs derived from feces at the peak of treatment *in vivo* are most distinct from the composition of their inoculum. Weighted Unifrac distance of the 7^th^ passage in BHI of each SIC to their fecal inoculum. Symbols and line are as in (B); *n*=12. D) SIC diversity correlates with inoculum diversity in most media. Richness (number of ASVs in rarefied data) was compared for the 7^th^ passage of each SIC and the corresponding fecal inocula and separated and colored/shaped as in (A). *R* and *p* were calculated from all samples in a given medium; *n*=48. E) SIC yield scales with diversity. Mean OD of SICs after 20 h of growth in BHI passages 3-7 increases with increasing richness (number of ASVs in rarefied data) in passage 7. *R*, Pearson correlation coefficient; *n*=48. F) Maximum growth rate is correlated with the abundance of fast-growing families. Maximum growth rate was calculated from growth curves during the 7^th^ passage and BHI-passaged SICs were classified based on the relative abundance of Enterobacteriaceae and Enterococcaceae (*f*_Entero_). Black circles, mean maximum growth rate for each group (*n*=11 for *f*_Entero_ < 20% and *n*=37 for *f*_Entero_ > 20%); error bars, SD. Individual data points are plotted in gray. *p*-value is from a Student’s two-sided *t*-test between the two groups. G) SIC yield scales with the presence of fast-growing families. OD_20 h_ is as in (E). SICs with >20% Enterococcaceae but <20% Enterobacteriaceae are labeled as *f_Enterococcaceae_*>20%. Black circles, mean maximum growth rate for each group (*n*=5 for *f*_Entero_ < 20%, *n*=15 for *f_Enterococcaceae_*>20%, and *n*=8 for *f*_Entero_ > 20%); error bars, SD. Individual data points are plotted in gray. *p*-values are from a Student’s two-sided *t*-test between each pairwise comparison. H-I) Changes in SIC composition during passaging are reflected in altered SIC growth. Percentage of Enterococcaceae (H) and maximum growth rate (I) for three technical replicates (i-iii) of the SIC derived from SD mouse 2 during residual treatment passaged in YCFA. The increase in maximum growth rate and blooming of the Enterococcaceae occur concurrently in two of the replicates.

### SIC and fecal-sample richness are highly correlated due to maintenance of abundant species and within-family replacement

The presence or absence of multiple families in the first principal coordinate, and the observation that this coordinate mainly separated SICs by their inocula (Fig. 2A), suggested that SIC richness depends on the richness of the inoculum. Indeed, the first principal coordinate was correlated with inoculum richness (Fig. S3C), and the richness of SICs grown in BHI, TYG, and GAM was highly correlated with that of the inoculum, with some SICs having >50 detectable ASVs (Fig. 2D). These SICs showed similar evenness (relative abundance homogeneity) as their inocula (Fig. S4A). SICs grown in YCFA exhibited very low diversity and extreme evenness (Fig. 2D, S4A).

The total number of ASVs present in at least one inoculum was 158, and the total number of ASVs present in at least one SIC was 117. Nevertheless, only 79 of the 158 ASVs (50%) present in the inocula were present in at least one SIC, indicating emergence of some ASVs from below the limit of detection (Supplementary text). Most of the retained ASVs (68 of 79) were present in at least one BHI SIC. While fecal diversity was approximately maintained in SICs (Fig. 2D) and family-level composition was preserved for families conducive to culturing (Fig. 1D, 2A), passaging led to a reproducible decrease of some ASVs to below the limit of detection; this decrease was biased toward ASVs that started at lower abundance and was mostly balanced by emergent ASVs from the same family (Supplementary text). The only family to emerge above the limit of detection during passaging was the sole Gammaproteobacterial family (an Enterobacteriaceae). Nonetheless, the Enterobacteriaceae emerged almost exclusively in SICs derived from pre- and post-treatment inocula (Fig. S1M), indicating that this emergence was not due to contamination. Thus, both maintenance and within-family replacement contribute to SIC richness at steady state.

### SIC growth dynamics reflect compositional characteristics

We speculated that SIC characteristics such as richness and the presence or absence of key species may be reflected in the features of continuous growth curves. We first focused on BHI SICs since they harbored the highest diversity (Fig. 2D). We measured SIC optical densities (ODs) at 20 h (“yield”) because by that time all growth rates had become virtually zero. We averaged the last 4 passages for each condition and sample and found a range of mean final ODs that were highly correlated with SIC richness (*R*=0.92, *p*=6×10^-30^; Fig. 2E). Similar relationships held for TYG and GAM (Fig. S5A). Thus, the yield of these SICs was greater when more taxa are present, suggesting that in the appropriate medium, SICs with low diversity could support the growth of more taxa if they were present in the inoculum.

We next calculated the maximum growth rate (Methods) for each SIC and again observed that richness correlated with maximum growth rate (*R*=0.50, *p*=2.7×10^-4^, Fig. S5B). The distribution of maximum growth rates was bimodal, reaching as high as 1.7 h^-1^ (doubling time ∼25 min) (Fig. S5B), similar to the growth rates of fast-growing species such as *E*. *coli* in rich media (Baev et al., 2006). We hypothesized that the presence of those families would correlate with SIC maximum growth rate. The summed relative abundance of the two families (*f*_Entero_) was also bimodal and correlated with maximum growth rate (Fig. S5C). SICs with *f*_Entero_ > 20% had a significantly higher maximum growth rate (Fig. 2F). Yield (Methods) was also linked to the presence of Enterobacteriaceae and Enterococcaceae after 48 h of growth (Fig. 2G). These correlations were similar in TYG and GAM (Fig. S5D), indicating that the relationship between these families and growth rate is likely medium-independent.

In YCFA, there was no correlation between richness and final OD (Fig. S5A), probably due to the overall low diversity of these SICs. Nonetheless, the relative abundance of fast growers was still linked to maximum growth rate (Fig. S5D). Some YCFA technical replicates that diverged in composition during passaging exhibited blooming of an *Enterococcus* species in the sixth passage in two of three replicates (Fig. S1Jiii). This bloom coincided with a change in maximum growth rate (Fig. 2H,I), suggesting that growth features can also be used to dynamically detect major changes in composition *in vitro*.

### The population growth behavior of SICs can be partially reconstituted using isolates

Ideally, our top-down approach to SIC cultivation could be complemented with bottom-up reconstitution using isolates. We focused on an SIC cultured in BHI from a MD mouse pre-treatment (Pre-MD) inoculum (parent SIC, Fig. 1D,3A) based on its high diversity (>50 ASVs, Fig. 2D). We obtained 15 distinct isolates by plating and analyzing 192 colonies (Fig. 3A, Methods). While these isolates represented <30% of the total number of ASVs in the parent SIC, they accounted for 69% of its total abundance (Fig. 3B). Our isolates included the sole Enterococcaceae, Porphyromonadaceae, and Enterobacteriaceae ASVs present at >0.1% in the parent SIC (Fig. 3B).

**Figure 3:**
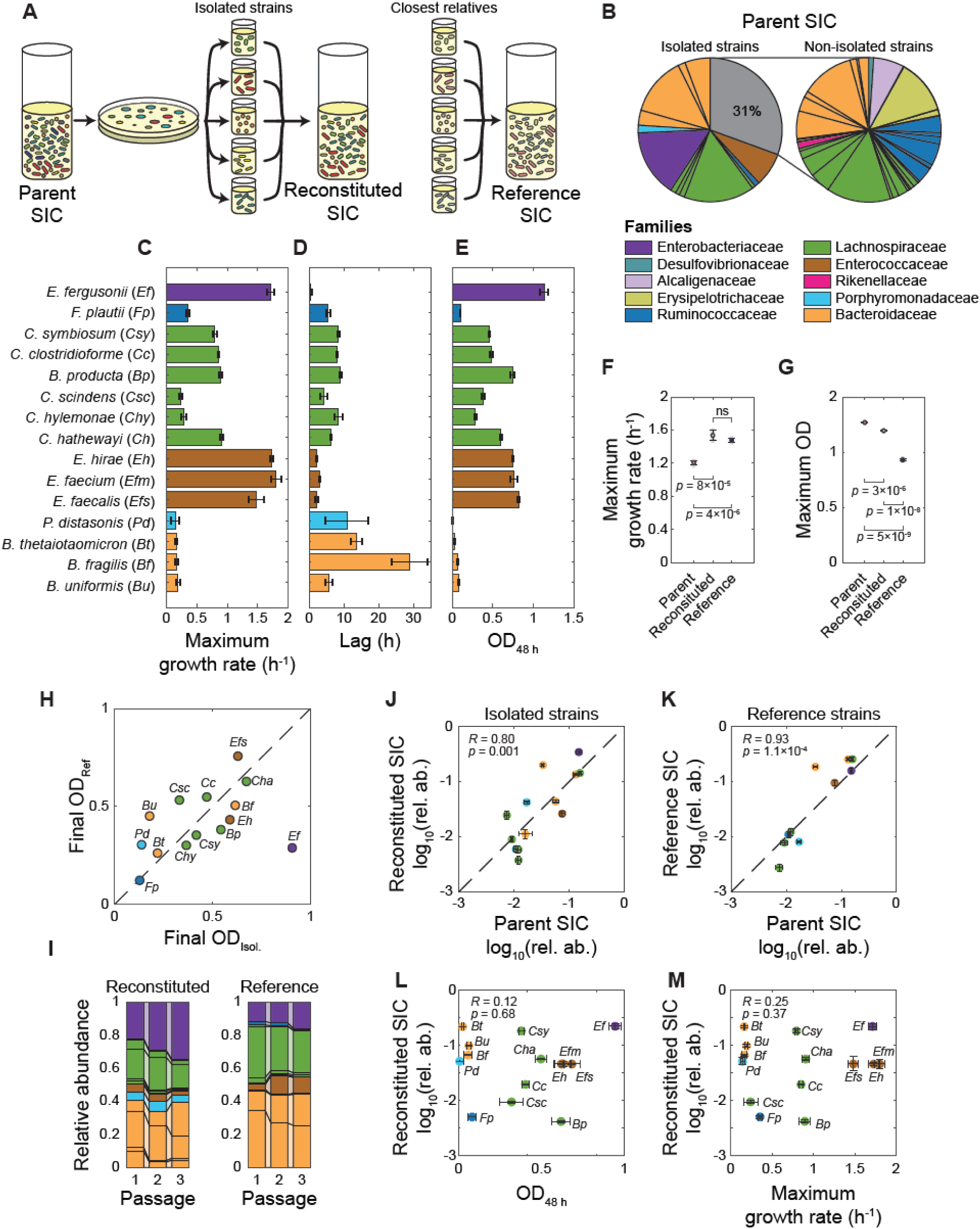
Community growth can be partially reconstituted from isolates. A) Experimental setup. Fifteen strains representing distinct species were isolated from an SIC passaged in BHI derived from an MD mouse pre-treatment fecal sample (parent SIC). The 15 strains, or the closest relative reference strains, were mixed and co-cultured (reconstituted and reference SIC, respectively). B) Isolated strains represent 69% of the composition of the parent SIC. Left: relative abundances of the 15 ASVs in the parent SIC. Right: normalized relative abundances of the non-isolated ASVs. Multiple strains map to a single ASV. C-E) Isolates display a range of growth behaviors. Mean values of maximum growth rate (C), lag time (D), and OD after 48 h of growth (E) of the 15 isolates in (B). Error bars, standard deviations (SD); *n*=6. F-G) Parent SIC growth characteristics are partially recapitulated by reconstituted and reference SICs. Maximum growth rate (F) and OD after 48 h of growth (G) of the parent SIC in (B), and the reconsitituted and reference SICs. Colored circles and error bars represent mean values and SD; *n*=4. *p*-values are from a Student’s two-sided *t*-test between each pairwise comparison; ns: not significant. H) Yields (OD after 48 h of growth in BHI) of isolates in (C-E) and corresponding reference strains are mostly correlated. Isolate abbreviations are defined in (C). I) Relative abundances of ASVs in reconstituted (left) or reference (right) SICs during three passages in BHI. Multiple strains can be mapped to a single ASV. J,K) Reconstituted (J) and reference (K) SICs converge to compositions with relative abundances similar to that of the parent SIC. Data are mean log_10_(relative abundance) of ASVs, normalized to the total relative abundance of those ASVs within the reconstituted or parent SIC. Error bars, SD; *n*=4. *R*, Pearson correlation coefficient. L,M) Isolate growth behaviors do not correlate with their abundances in reconstituted SIC. Mean log_10_(relative abundance) of ASVs corresponding to isolates in the 3rd passage of reconstituted SIC is uncorrelated with the OD of the isolates after 48 h of growth (L) or maximum growth rate of those strains in isolation (M). Error bars, SD; *n*=4 for relative abundances, *n*=6 for OD and maximum growth rate. *R*, Pearson correlation coefficient. Where multiple strains map to a single ASV, the relative abundance of the ASV value is used for all strains that map to that ASV.

To test whether these 15 isolates were sufficient to recapitulate the growth and composition of the parent SIC, we examined the growth behaviors of our parent SIC, as well as the isolates alone and in a reconstituted SIC (Fig. 3A). We grew each isolate from frozen stock in the medium from which it was isolated at 37 °C for 48 h. We then passaged the isolates individually in BHI and measured OD over time to quantify lag time, maximum growth rate, and yield. *Enterococcus* and *Escherichia* strains had higher maximum growth rates and shorter lag times than the other isolates (Fig. 3C,D), and were relatively abundant in the parent SIC (7.6±0.8% and 15.1±0.1%, respectively; *n*=3, Fig. 3B). While the initial phase of growth after subculturing is likely dominated by these fast-growing species, the parent SIC grew significantly slower than the three *Enterococcus* isolates (all of which were represented by the same ASV) and *Escherichia fergusonii* (Fig. S6A); the lag time of the parent SIC was higher than the shortest isolate lag time (*E. fergusonii*) (Fig. S6B). Isolate yield was more homogeneous (Fig. 3E). Although some of the poorest growers were *Bacteroides thetaiotaomicron* and *Parabacteroides distasonis* (final ODs 0.032±0.004 and 0.010±0.006, respectively, Fig. 3C), the corresponding ASVs ended up as 13.23±0.6% and 1.7±0.1% of the parent SIC, respectively, suggesting that their growth is not greatly hampered by the initial expansion of other species.

We built a reconstituted SIC (Fig. 3A) by growing the 15 isolates independently to saturation, diluting them to OD=0.001 in BHI, and mixing them in equal volumes. To test whether other strains slowed the growth of *Enterococcus* spp. and *E. fergusonii* in the parent SIC, we monitored growth at 37 °C for 48 h and then passaged this reconstituted SIC twice more. The reconstituted SIC had a higher maximum growth rate but the same lag time as its parent SIC in the third passage (Fig. 3F, S6C). Thus, fast-growing species likely expand initially in the absence of competition for nutrients, reshaping the niche landscape, after which slower-growing species fill unoccupied or new niches and enforce physiological states on the fast-growing species that make them unable to grow as quickly in subsequent passages despite their higher relative abundance. Supporting this notion, the yield of the parent SIC was higher than that of the reconstituted SIC in the third passage (Fig. 3G); this increase was realized during entry to stationary phase (Fig. S6D), indicating that the other, unisolated species were slow growers that occupy niches not filled by the isolates. Both sets of species contribute to yield and further depress maximum growth rate.

Are the growth properties of reconstituted SICs general to the component species or specific to isolated strains? We built an SIC with the closest relatives of each of the 15 isolates that were available in reference strain collections (reference SIC, Fig. 3A) and monitored growth of the reference strains for one passage and of the reference SIC for three passages. The maximum growth rate of the isolates and their closest relative reference strains grown alone were highly correlated (Fig. S6E); the reference SIC had the same maximum growth rate (Fig. 3E) and a longer lag phase (Fig. S6C) than the reconstituted SIC. However, the reference SIC yield was substantially lower than that of the reconstituted SIC and the parent SIC (Fig. 3G), likely because *E. fergusonii* had the highest yield of all isolates but the corresponding reference strain exhibited a much lower yield in BHI (Fig. 3H). Hence, the ability of co-cultures to recapitulate a complex SIC can depend on community effects (growth rate) and metabolic properties of individual strains (yield).

To test how much the composition of reconstituted and reference SICs resembled that of the parent SIC, we performed 16S amplicon sequencing (Methods). The reconstituted and reference SICs mostly stabilized within one passage (Fig. 3I). In general, the relative abundances of the ASVs in the reconstituted and reference SICs were highly correlated with those in the parent SIC (Fig. 3J,K), but not with isolate yields or maximum growth rates (Fig. 3L,M). The family-level relative abundances in reconstituted and reference SICs were correlated (Fig. S6F). Notably, even though *E. fergusonii* and *Enterococcus* spp. can grow more quickly and more than most other isolates (Fig. S6E), their overall contributions to co-culture yield were limited by the other species. Moreover, the similarly high abundances of the 3 *Bacteroides* species (despite all but *B. fragilis* being poor growers in isolation in unsupplemented BHI, Fig. S6E) suggests that some *Bacteroides* rely on other organisms for optimal growth. Overall, co-culturing these 15 species seems to reproduce much of the metabolic web responsible for establishing their abundances in the parent SIC.

### Ciprofloxacin alters SICs in a manner generally consistent with isolate sensitivities

We queried the responses of our SICs to ciprofloxacin. We selected an SIC derived from an inoculum that had not been exposed to the drug *in vivo* (pre-treatment from SD mouse, Pre-SD, Fig. 4A). We grew a frozen stock in BHI for 48 h over two rounds of passaging and then diluted 1:200 into a 96-well plate in duplicate with increasing ciprofloxacin concentrations (Fig. 4Ai). The maximum growth rate of the SICs decreased monotonically with drug concentration (Fig. 4B) and the collective minimum inhibitory concentration (MIC) for the SIC was >32 μg/mL (Fig. 4B). The final OD of the Pre-SD SIC exhibited at least two notable decreases, at the lowest and highest concentrations (Fig. 4B); our findings (Fig. 2A) suggest that these decreases are due to drops in diversity. To test the effects of long-term exposure to ciprofloxacin, we passaged the Pre-SD SICs in BHI with the ciprofloxacin concentrations to which they were exposed in the first passage for two more *in vitro* passages (Fig. 4Aii,iii). For the highest concentrations, the maximum growth rate increased moderately after the first passage (Fig. 4C). The increasing growth rate upon additional passages in antibiotic suggest that the structure of the SIC changes after *in vitro* treatment, potentially due to selection of resistant subpopulations.

**Figure 4:**
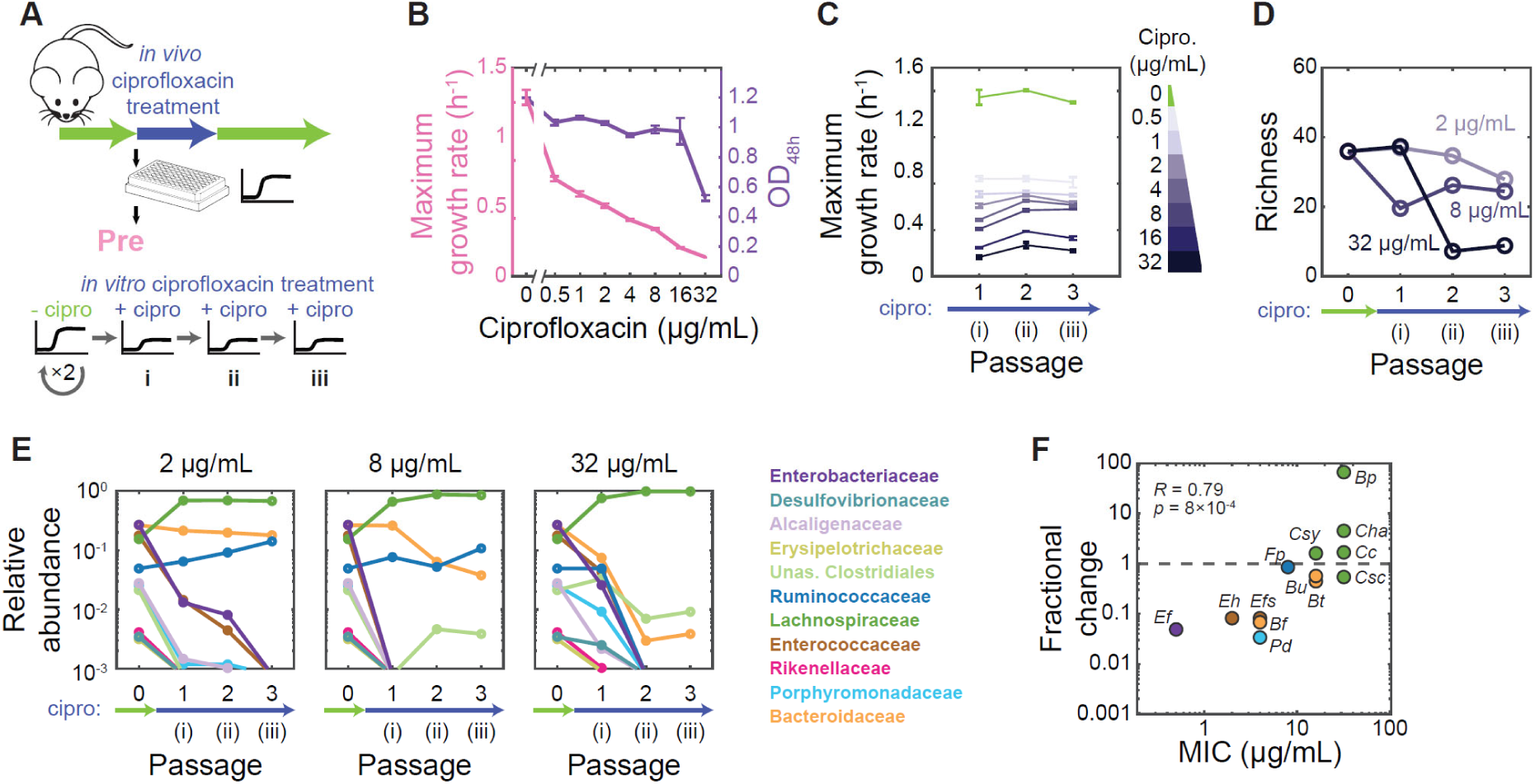
SIC response to ciprofloxacin can be predicted from exposure history and individual member sensitivities. A) Experimental setup. An SIC passaged in BHI from pre-treatment humanized mouse fecal inoculum (Pre-SD) was revived after freezing and passaged twice in BHI. The SIC was then passaged in ciprofloxacin three times. B) Pre-SD SIC yield and growth rate decrease with increasing concentrations of ciprofloxacin. OD was measured after 48 h of growth with ciprofloxacin. Lines, means of triplicate growth curves; error bars, standard deviations (SD). C) Pre-SD SIC shows little adaptation to ciprofloxacin. Maximum growth rate decreases or remains approximately constant across three passages in ciprofloxacin. Symbols are as in (B). D) Richness (number of ASVs in rarefied data) of Pre-SD SIC decreases during ciprofloxacin. Data are means of two technical replicates. E) Ciprofloxacin treatment *in vitro* selects for a few families. Family-level composition of the Pre community across passages in ciprofloxacin. The limit of detection was estimated at 10^-3^. Data are means of two technical replicates. F) Growth of ASVs in the Pre-SD SIC under ciprofloxacin treatment can be predicted by their sensitivity in isolation. Fractional-abundance changes of ASVs corresponding to the 15 isolates in Fig. 3A were computed after one passage in 2 μg/mL ciprofloxacin. MICs were calculated during growth in isolation in BHI (Table S1).

To assess compositional changes in the Pre-SD SIC upon treatment, we performed 16S sequencing after 48 h of treatment with 2, 8, and 32 μg/mL ciprofloxacin in each of the three consecutive passages (Fig. 4Ai,ii,iii). At 2 μg/mL, diversity of the Pre-SD SIC decreased relative to the untreated SIC in the first passage (Fig. 4D). At 32 μg/mL, the diversity of the Pre-SD SIC declined to levels lower than those of SICs derived from residual-treatment inocula of the same SD mouse in the absence of ciprofloxacin (Fig. 2D, 4D), suggesting that the 2-32 μg/mL range encompasses those experienced by the microbiota *in vivo*. For all doses, many families in the Pre-SD SIC decreased to undetectable levels, including the abundant Enterobacteriaceae and Enterococcaceae (Fig. 4E), consistent with the drop in maximum OD relative to the untreated SIC (Fig. 2G, 4B). Members of the Lachnospiraceae, Ruminococcaceae, and Bacteroidaceae persisted after three rounds in 2 and 8 μg/mL ciprofloxacin (Fig. 4E). At 32 μg/mL, the Ruminococcaceae were undetectable by the second passage and the Bacteroidaceae decreased to very low levels (Fig. 4E), likely explaining the large decrease in maximum OD relative to 16 μg/mL (Fig. 4B).

Next, we examined changes at the ASV level to assess within-family dynamics. In the Pre-SD SIC, antibiotic pressure can select for strains that emerge from below the detection limit due to the decrease of other, more sensitive species: at 2 μg/mL the Lachnospiraceae increased from 12 to 16 detectable ASVs while the Bacteroidaceae decreased from 12 to 4 detectable ASVs. At every concentration, the changes were highly reproducible between replicates (more than 89% and 69% of ASVs that became undetectable or detectable, respectively, did so in both replicates). To test whether the observed changes were explained by the sensitivities of each family to ciprofloxacin, we measured the MIC of the 15 strains isolated from a Pre-MD SIC (Fig. 3A, Table S1). The isolate MICs were highly correlated with the fractional change in abundance upon 2 μg/mL ciprofloxacin treatment of the Pre-SD SIC (Fig. 4F). These data show that the compositional changes observed in SICs can largely be predicted by individual sensitivities in isolation.

### SICs can predict *in vivo* compositional changes due to ciprofloxacin

While a drug concentration can be measured in fecal samples, it is usually undetermined along the gastrointestinal tract and potentially heterogeneous in space and time. Other variables are currently impossible to measure and control, such as environmental reservoirs that could play a role in bacterial migration and repopulation. Our SICs provide the opportunity to study antibiotic perturbations in a precisely controlled and high-throughput manner, to assess similarity to *in vivo* behavior, and to interpret changes *in vivo* mechanistically.

Humanized mice treated with ciprofloxacin harbored high levels of Lachnospiraceae and Ruminococcaceae at the peak of treatment (Fig. S1A), in agreement with the selection of these families in SICs (Fig. 4E). Of the families for which we obtained MICs (Table S1), the highest MIC was observed in the Lachnospiraceae, and this family remained high or increased in relative abundance upon residual treatment *in vivo* (Fig. 5A). Most sensitive families (MIC <32 μg/mL) decreased in abundance (Fig. 5A, Fig. S1M); the lone exception was the Bacteroidaceae, which increased in three of four mice (Fig. 5A), consistent with our previous observation that a highly resistant *Bacteroides vulgatus* strain (MIC >512 μg/mL) can be selected *in vivo* (Ng et al., 2019). Again, these results suggest that the effective concentration *in vivo* is between 2 and 32 μg/mL. These similarities suggest that ciprofloxacin sensitivities are generally similar *in vitro* and *in vivo*, and that individual sensitivities largely dictate the initial dynamics of SIC composition *in vivo*.

**Figure 5:**
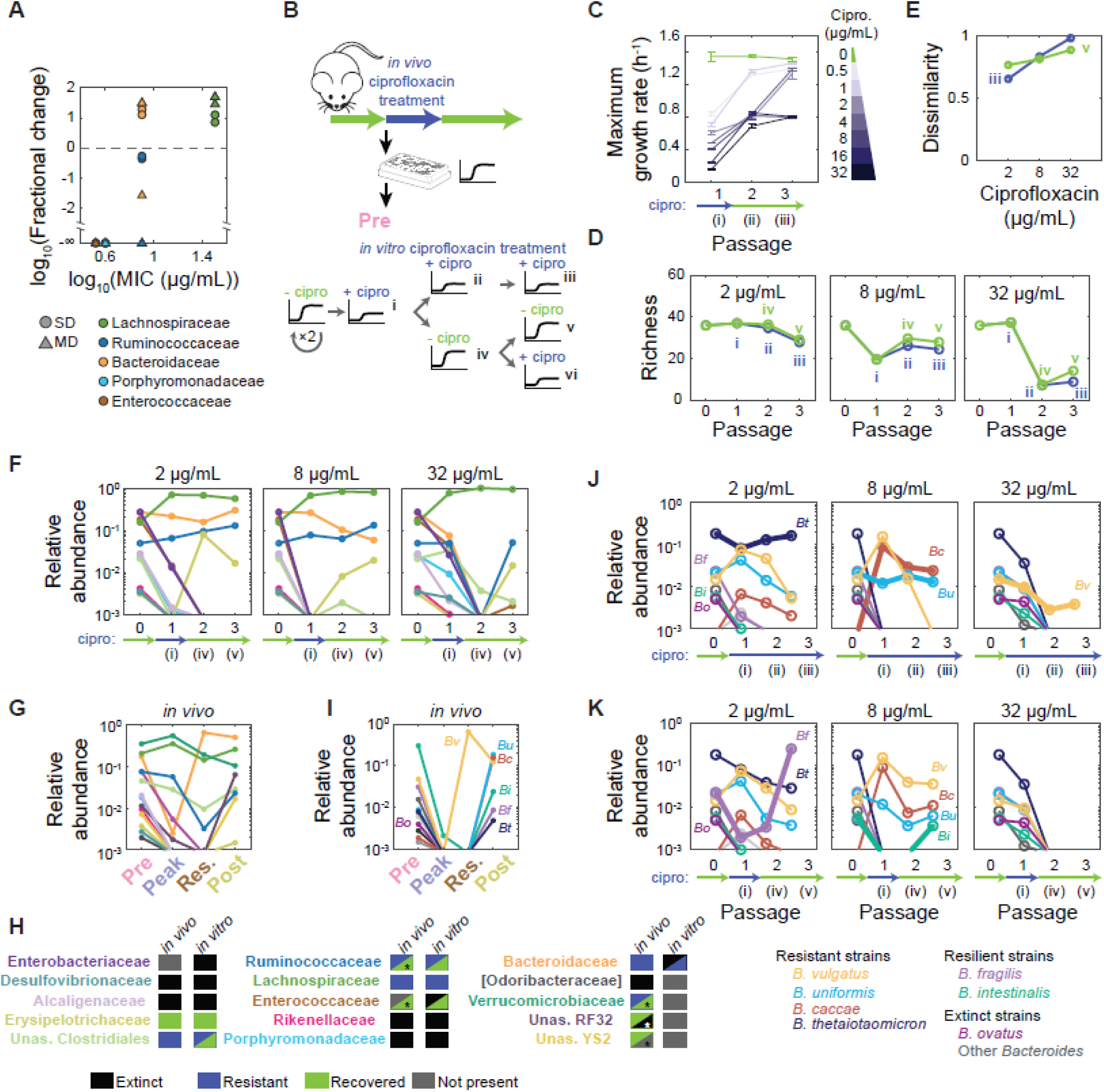
Ciprofloxacin treatment *in vitro* reproduces features of *in vivo* dynamics at multiple taxonomic levels. A) Changes in family-level abundances during ciprofloxacin treatment *in vivo* can be predicted by strain-level sensitivities *in vitro*. Fractional changes were computed as the ratio of residual and pre-treatment time-point abundances *in vivo*. MIC is the mean value across strains in a given family determined from growth in isolation in BHI (Table S1). B) Experimental setup mimicking *in vivo* transient treatment. An SIC passaged in BHI from pre-treatment humanized mouse fecal inoculum (Pre-SD) was revived after freezing and passaged twice in BHI. The SIC was then passaged in ciprofloxacin three times (i,ii,iii); in ciprofloxacin once and then twice without the drug (i,iv,v); or re-exposed to the drug after one round of recovery (i,iv,vi). C) Pre-SD SIC growth rate can recover to levels similar to values before treatment at low concentrations. Maximum growth rate was calculated across ciprofloxacin concentrations during one round of antibiotic treatment and two rounds of recovery. Lines, mean of three technical replicates; error bars, standard deviations (SD). D) Richness of Pre-SD SIC recovered only partially after removal of ciprofloxacin (green) versus continuous exposure (blue, same data as Fig. 4D). Data are the mean of two replicates. E) The Pre-SD SIC was further remodeled after removal of low-dose antibiotic treatment. Bray-Curtis dissimilarity index of SICs was calculated after the third passage in ciprofloxacin (blue) or after two rounds of recovery (green) with respect to their untreated composition. Data are the mean of two technical replicates. F) Some families, like the Erysipelotrichaceae and Ruminococcaceae, can recover after transient ciprofloxacin treatment. Data are the mean log_10_(relative abundance) of two replicates during one round of ciprofloxacin treatment and two rounds of recovery at the family level. G) Ciprofloxacin-induced changes to family-level abundances in SD mice *in vivo* resemble changes *in vitro* in (F). Data are the mean log_10_(relative abundance) of two mice at the family level. H) Most families display similar qualitative abundance changes *in vivo* and *in vitro*. Split boxes denote either different responses across diets *in vivo* (asterisk denotes the response in MD mice, Fig. S9E) or across ciprofloxacin doses *in vitro*. I) *Bacteroides* dynamics *in vivo* consist of *B. vulgatus* dominance during treatment and the recovery of several species after treatment. Data are the mean log_10_(relative abundance) of two SD mice at the ASV level: *Bacteroides ovatus* (*Bo*), *B*. *vulgatus* (*Bv*), *B*. *uniformis* (*Bu*), *B*. *caccae* (*Bc*), *B*. *intestinalis* (*Bi*), *B*. *fragilis* (*Bf*), and *B*. *thetaiotaomicron* (*Bt*). Other *Bacteroides* are shown in shades of grey. J,K) *Bacteroides* survival and recovery *in vivo* can be respectively explained by resistance and resilience characteristics of the Pre-SD SIC *in vitro* during continuous treatment (J) or one round of treatment followed by two rounds of recovery (K). Data are the mean log_10_(relative abundance) of two replicates during one round of ciprofloxacin treatment and two rounds of recovery at the ASV level. Thick lines highlight *Bacteroides* ASVs that show resistance (J) or resilience (K) at the highest concentration that they display the behavior.

*In vivo*, the microbiota undergoes a complex recovery that likely involves differential susceptibilities, intrinsic recovery capacities, the emergence of resistant mutants, and metabolic interactions (Ng et al., 2019). To investigate the response of SICs in conditions resembling recovery from treatment *in vivo*, we diluted the Pre-SD SIC into plates without ciprofloxacin after one round in ciprofloxacin (Fig. 5Bi,iv,v). The Pre-SD SIC exhibited increases in maximum growth rate after one round of treatment at all doses (Fig. 5C); at low doses, growth rate rebounded to values similar to those before treatment, while at >8 μg/mL it only partially recovered (Fig. 5C). All the doses at which maximum growth rate increased back to similar levels to those before treatment showed recovery of the Enterococcaceae (Fig. S7A), in line with the correlation of the presence of this family and maximum growth rate (Fig. 2B) and the MIC of *Enterococcus* isolates (2-4 μg/mL, Table S1). Removal of the drug slightly increased diversity in Pre-SD SICs (Fig. 5D), indicating that the recovery of certain species drives the partial recovery of growth rate. Thus, diversity recovery *in vitro* and *in vivo* (Fig. S1B) can be partially explained by re-emergence of ASVs that became undetectable during treatment.

We next computed the extent of compositional disruption during treatment and the degree of recovery after treatment based on the Bray-Curtis dissimilarity index. Ciprofloxacin dose determined the dissimilarity between the Pre-SD SIC before treatment and after three passages in ciprofloxacin (Fig. 5E). The compositions of the treated SICs were highly reproducible at the ASV level, especially at low concentrations (Fig. S7B). After removing the drug from the highest-concentration conditions, the SICs became more similar to, although still differed substantially from, the untreated state (Fig. 5E). Interestingly, at lower concentrations the dissimilarity increased after removal of the drug (Fig. 5E) due to the survival and expansion of taxa that were present at very low abundance before ciprofloxacin treatment. This non-monotonic dose-response is reminiscent of the better recovery of the microbiota in mice during high-dose compared with low-dose osmotic diarrhea (Tropini et al., 2018). SICs hence adopt a new steady state after ciprofloxacin exposure and recovery, as we previously observed *in vivo* (Ng et al., 2019).

The small increase in SIC diversity after ciprofloxacin removal (Fig. 5D) was associated with the recovery of Erysipelotrichaceae, Ruminococcaceae, and Enterococcaceae members from below the limit of detection (Fig. 5F). The Erysipelotrichaceae family recovered from one treatment at all concentrations (Fig. 5F), consistent with its recovery *in vivo* post-treatment (Fig. S1A). A second treatment after one passage without the drug (Fig. 5Bvi) again made the Erysipelotrichaceae undetectable (Fig. S7C), indicating that sensitivity to ciprofloxacin was the same before and after one treatment-recovery cycle. The other families did not recover from undetectable levels after removing the drug (Fig. 5F). Of the families that became undetectable, the only one to exhibit different recovery characteristics *in vitro* and *in vivo* (Fig. 5F,G,H, S7D) was the Bacteroidaceae, which were undetectable at 32 μg/mL ciprofloxacin *in vitro* but persisted *in vivo*. Taken together, these results show that *in vitro* perturbations can be used as a proxy for many aspects of *in vivo* dynamics.

### Bacteroides recovery in vitro is due to a combination of resistance and resilience

The unexpected recovery of the Bacteroidaceae *in vivo* motivated us to probe the dynamics of these ASVs. *In vivo*, although most *Bacteroides* ASVs were undetectable at the peak and residual time points, most reappeared post-treatment (Fig. 5I, S7E). Only *B. vulgatus* in relative abundance to become dominant during residual treatment (Fig. 5I, S7E); this strain was likely highly resistant to ciprofloxacin based on our assessment of *Bacteroides* isolates from the residual fecal sample (Ng et al., 2019). Consistent with this observation, *B. vulgatus* was the only Bacteroidaceae ASV that was present *in vitro* after 3 passages with 32 μg/mL ciprofloxacin, reaching a stable abundance (Fig. 5J). *B. caccae* and *B. uniformis* were present after 3 passages at 8 μg/mL, and *B. thetaiotaomicron* was present after 3 passages at 2 μg/mL (Fig. 5J); all three species recovered *in vivo* post-treatment (Fig. 5I). Other *Bacteroides* spp., such as *B. fragilis*, became undetectable after one passage at 2 μg/mL (Fig. 5J). These trends are consistent with the higher MICs of *B. uniformis* and *B. thetaiotaomicron* compared to *B. fragilis* (Table S1). Thus, sensitivities of isolates are predictive of *Bacteroides* resistance during continuous ciprofloxacin treatment in SICs.

Like *B. fragilis*, *B. intestinalis* was undetectable by the second passage even at the lowest ciprofloxacin concentration tested *in vitro* (Fig. 5J). Nonetheless, both species recovered *in vivo* in SD mice (Fig. 5I). Remarkably, these species also re-emerged *in vitro* when ciprofloxacin-treated SICs were diluted into medium without the drug (Fig. 5K). *B. fragilis* increased in relative abundance and became dominant after treatment with 2 μg/mL (Fig. 5K), while *B. intestinalis* was detected only after 8 μg/mL (Fig. 5K), suggesting that competition with other species or taxa at the lower concentration drove *B. intestinalis* to undetectable levels. After treatment with 32 μg/mL, no *Bacteroides* species recovered (Fig. 5K). We refer to the ability of these strains to recover from undetectable levels during treatment as “resilience”. Interestingly, *B. vulgatus* was only detected when the SIC was continuously exposed (Fig. 5J) and not after ciprofloxacin was removed (Fig. 5K), suggesting that the recovery of other taxa can drive resistant members to undetectable levels. Species that did not recover *in vivo*, such as *B. ovatus*, did not recover *in vitro* (Fig. 5I,K). These data demonstrate that fecal-derived SICs can recapitulate species-level dynamics observed *in vivo* and can distinguish between resistance and resilience. Our observations of species resilience within complex SICs also show that re-seeding from environmental reservoirs is not necessary for species to recover after antibiotic treatment.

### Exposure to ciprofloxacin in vivo leads to functional differences in vitro

The compositions of SICs derived during and after antibiotic treatment resembled the compositions of the inocula (Fig. 2A,C). Thus, we next asked whether the SICs retained functions observed *in vivo*: decreased resistance to colonization by the pathogen *Salmonella enterica* serovar Typhimurium (Barthel et al., 2003; Lawley et al., 2008) and increased resilience to a second round of treatment (Ng et al., 2019).

First, we asked whether exposure to ciprofloxacin led to compositional changes that reduce colonization resistance in the absence of the drug *in vitro*. We grew SICs derived from SD mice inocula gathered before, during the peak of, during residual, and after treatment *in vivo* (Pre-SD, Peak-SD, Res-SD, and Post-SD, respectively, Fig. 6A) in BHI. We challenged these SICs with *S.* Typhimurium for 48 h in BHI, diluted them, spotted them onto LB-streptomycin agar plates, and incubated them in aerobic conditions to select for *S.* Typhimurium. Pre-SD SICs had >10-fold less *S.* Typhimurium than communities derived from feces of ciprofloxacin-treated mice (Fig. 6B, S8A). Single-cell quantification of fluorescently tagged *S.* Typhimurium revealed an even larger difference in colonization efficiency between the Pre-SD and Res-SD SICs (Fig. 6C, S8B). Thus, changes in composition driven by antibiotic treatment *in vivo* lead to differences in SICs that mimic the resilience of the microbiota to pathogen invasion *in vivo*.

**Figure 6:**
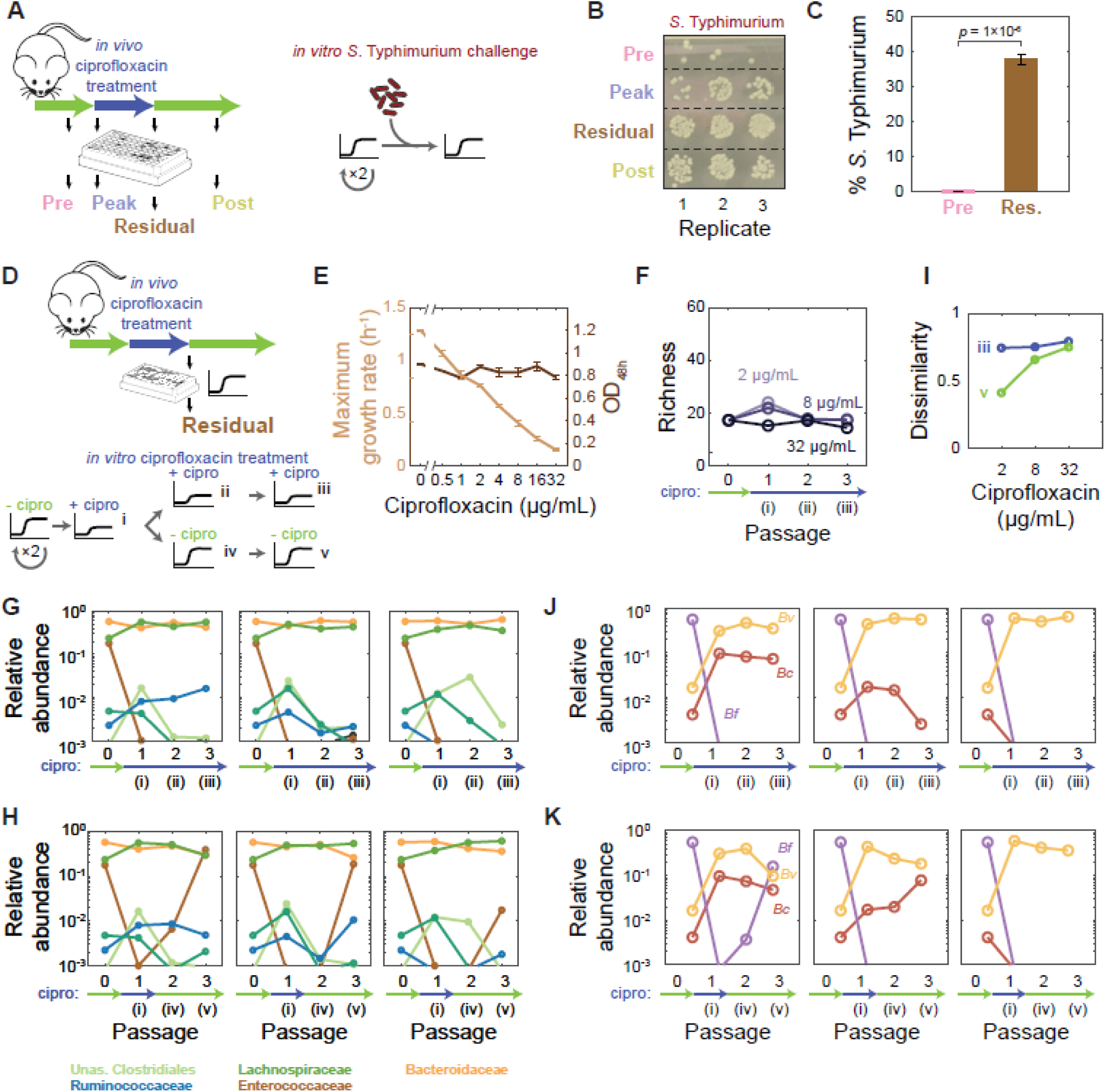
Pre-exposure of the gut microbiota to ciprofloxacin *in vivo* results in functional differences *in vitro*. A) Experimental setup for *in vitro* challenge with *S*. Typhimurium. SICs passaged in BHI from pre-, peak, residual, and post-treatment humanized mouse fecal inocula were revived after freezing and passaged twice in BHI. SICs were mixed with *S*. Typhimurium and *S*. Typhimurium levels were quantified after 48 h of growth. B) SICs derived from mice treated with ciprofloxacin are more susceptible to *S*. Typhimurium. Colonies of *S*. Typhimurium SL1344 after 48 h of growth with SICs diluted 1:10^4^ and grown aerobically on LB+streptomycin. C) Single-cell quantification of mCherry-tagged *S*. Typhimurium 14028s after co-culture with SICs derived from pre- and residual-treatment mice fecal inocula. *p*-value is from a Student’s two-sided *t*-test; *n*=3. D) Experimental setup for *in vitro* antibiotic treatment of a pre-exposed SIC. An SIC passaged in BHI from residual treatment humanized mouse fecal inoculum (Res-SD) was revived after freezing and passaged twice in BHI. The SIC was passaged in ciprofloxacin three times (i,ii,iii) or in ciprofloxacin once and then twice without the drug (i,iv,v). E) In contrast to the Pre-SD SIC (Fig. 4B), the yield and of Res-SD SIC is virtually unaffected by ciprofloxacin, although the growth rate decreases with concentration. OD was measured after 48 h of growth with ciprofloxacin. Lines, means of triplicate growth curves; error bars, standard deviations (SD). F) Richness (number of ASVs in rarefied data) of Pre-SD SIC remains constant during continuous ciprofloxacin treatment. Data are means of two technical replicates. G) Bacteroidaceae and Lachnospiraceae dominate during three rounds of ciprofloxacin treatment of the Res-SD SIC. Data are the mean of family-level abundances across two technical replicates. H) Enterococcaceae can recover after one round of ciprofloxacin treatment in the Res-SD SIC. Data are the mean of two replicates. I) Res-SD SIC diversity can recover after ciprofloxacin removal. The Bray-Curtis dissimilarity index of SICs at the third passage in ciprofloxacin (blue) or after two rounds of recovery (green), compared to their untreated compositions. Data are the mean of two replicates. G,H) Bacteroidaceae are remodeled in the Res-SD SIC during and after ciprofloxacin treatment, with *B. vulgatus* as the only member to generally survive. Relative abundances of *Bacteroides* species in the Res-SD SIC during continuous ciprofloxacin treatment (G) or after one round of treatment followed by two rounds of recovery (H).

Second, we asked whether the pre-exposed, low-diversity Res-SD SIC showed higher resilience to ciprofloxacin *in vitro* than the Pre-SD SIC (Fig. 6D). We grew the SICs in BHI with ciprofloxacin continuously (Fig. 6Di,ii,ii) or transiently (Fig. 6Di,iv,v) as above. The Res-SD SIC was less sensitive than the Pre-SD SIC in terms of growth rate (Fig. 4B, 6E), and the final OD was largely unaffected (Fig. 6E).

We next asked whether the robustness of Res-SD SIC growth behaviors (Fig. 6E) during ciprofloxacin treatment could be explained by particular compositional differences from the Pre-SD SIC. The Res-SD SIC displayed responses distinct from those of the Pre-SD SIC, maintaining diversity even at the highest ciprofloxacin concentrations (Fig. 4C, 6F). At all concentrations, Bacteroidaceae and Lachnospiraceae dominated even after three passages (Fig. 6G), in contrast to the elimination of most Bacteroidaceae in the Pre-SD SIC (Fig. 4E). The most abundant family that was subsequently lost from Res-SD SIC was the Enterococcaceae (Fig. 6G); the Ruminococcaceae and Verrucomicrobiaceae were lost at 32 and 2 μg/mL ciprofloxacin, respectively (Fig. 6G). These losses were counterbalanced by emergent ASVs within the resistant families. As with the Pre-SD inocula, composition was quantitatively similar at the ASV level between the two replicates at low ciprofloxacin concentrations (Fig. S7G). At 32 μg/mL, the two replicates were more similar to each other than to the Pre-SD SIC after treatment (Fig. S7D), suggesting that pre-exposure to the drug in mice decreases the stochastic nature of species elimination upon subsequent exposure *in vitro*.

The compositions of the Res-SD SIC before treatment and after two recovery passages were highly similar at low doses (Fig. 6H). For all doses, the dissimilarity index of the post-recovery Res-SD SIC reverted toward its original state (Fig. 6I); by contrast, at 2 μg/mL ciprofloxacin, the Pre-SD SIC was further remodeled during recovery (Fig. 5E). These data imply that the members of the Res-SD SIC are more resistant to the drug than members of the Pre-SD SIC, in line with their pre-selection *in vivo*.

Both *in vitro* for the Pre-SD SIC and *in vivo*, we observed selection of *B. vulgatus* upon treatment with ciprofloxacin (Fig. 5I,J). However, in the absence of ciprofloxacin, *B. fragilis* dominated the Bacteroidaceae of the Res-SD SIC (Fig. 6J), consistent with this species’ ability to recover and dominate the Pre-SD SIC after ciprofloxacin removal (Fig. 5K), while *B. vulgatus* and *B. caccae* were at low relative abundances (Fig. 6J). *B. fragilis* became undetectable during Res-SD treatment regardless of concentration, as did *B. caccae* at 32 μg/mL (Fig. 6J). *B. vulgatus* accounted for the totality of the Bacteroidaceae at 32 μg/mL (∼20% of the SIC; Fig. 6G,J). At 2 μg/mL, the *B. fragilis* ASV re-emerged after two recovery passages in one replicate at the expense of *B. vulgatus* (Fig. 6K), similar to the replacement of the resistant *B. vulgatus in vivo* by other *Bacteroides* (Fig. 5I). Thus, SICs derived from residual inocula are generally more resilient to ciprofloxacin and exhibited *Bacteroides* dynamics similar to *in vivo* treatment. These data suggest that the initial treatment culls the community down to a resistant core, leading to a community that is stable under further treatment.

### Diverse, stable SICs can be generated directly from human fecal samples

Our experiments consistently showed that passaging of fecal samples from humanized mice can lead to stable SICs preserving many of the taxa from the original sample. What about feces sampled directly from humans? Three human stools from anonymous donors were more diverse than the humanized mice fecal samples (Fig. 2D,S1B), with 115-219 ASVs. We passaged the samples in BHI and observed trends of rapid convergence to a stable steady state and high family-level diversity similar to SICs derived from humanized mice (Fig. 7A); SIC composition stabilized in 3.1±0.3 passages, and the mean richness after the 8^th^ passage was 46.4±2.4 ASVs (*n*=3).

**Figure 7:**
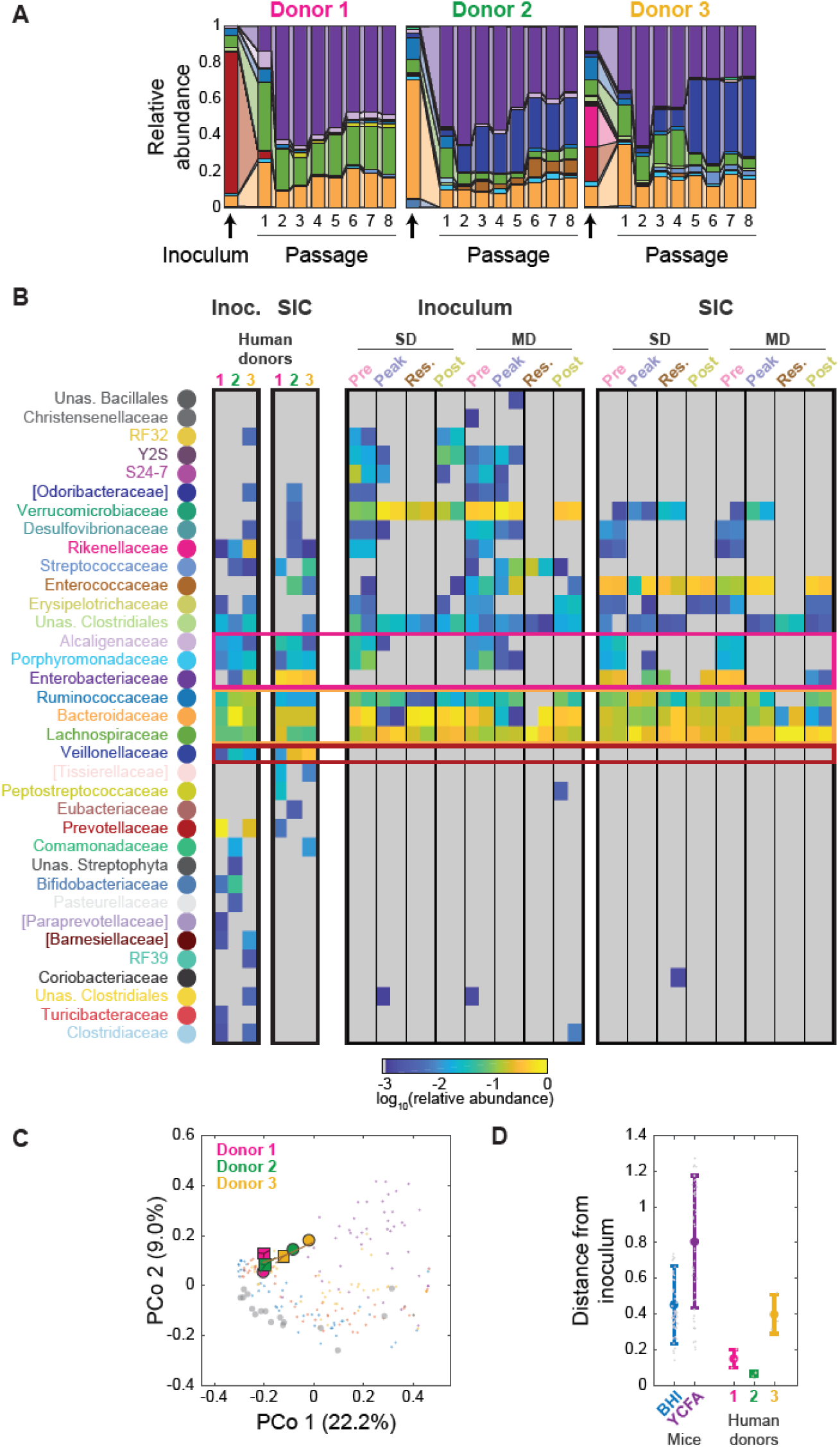
Batch culturing of human stool leads to stable and complex SICs. A) *In vitro* passaging of stool samples from three healthy human donors in BHI produced stable, complex SICs. Family-level composition; each passage was 48 h. Colors corresponding to individual families are as in (B). B) *In vitro* passaging of stool samples leads to SICs that maintain many families from the fecal microbiota. Mean relative abundance of three technical replicates for each inoculum and corresponding 7^th^ passage SIC in (A) (left) and all humanized mice SICs (right). Only families with mean relative abundance >0.1% are shown; undetectable taxa are shown in grey. Boxes highlight families common to all SICs (orange), present in all human donor SICs but underrepresented in humanized mice SICs (pink), and exclusive to all human donor SICs (red). C) Passaging in BHI preserves the beta diversity of human stool inocula. PCoA was computed with unweighted Unifrac distances using all 192 passaged SICs inoculated with humanized mouse samples in Fig. 2A and the SICs in (A). Colors and shapes are as in (C). Symbols represent the centroid of three replicates; lines connect replicates to the centroid. SICs from humanized mice are shown in colors in smaller dots in the same colors as in Fig. 1G and the humanized mice fecal samples are shown in gray. D) Human stool-derived SICs are as similar to their inocula as humanized mice-derived SICs, based on weighted Unifrac distances between steady-state SICs and their fecal inocula. SICs derived from humanized mice passaged in BHI or YCFA are shown for comparison (data from Fig. 2B). Colored circles are the mean of SICs (*n*=48 for BHI and YCFA humanized mouse SICs and *n*=3 for BHI human donor SICs). Error bars, standard deviations. Individual data points are shown in gray.

Steady-state compositions of the passaged SICs were reproducible across technical replicates (Fig. S9A), but across donors, the SICs differed at the family level, with >50% of families being unique to one donor (Fig. 7A, S9B). Of the 103 unique ASVs across all human stool-derived SICs, only 9 were shared among SICs derived from all three donors (6 *Bacteroides* spp., 1 Lachnospiraceae, 1 Enterobacteriaceae, 1 Ruminococcaceae). Families that were shared among the three human stool-derived SICs were highly overlapping with those of the core microbiota from our other humanized mouse-derived passaged SICs (Fig. 7B, S9C). Moreover the relative abundances of these ASVs were similar in all human stool-derived SICs (Fig. S9B).

To compare to humanized mouse SICs, we performed a PCoA on humanized mouse and human stool SICs together, considering the presence or absence of ASVs (unweighted Unifrac distances, Fig. 7C). SICs from two stool samples had phylogenetic compositions (weighted Unifrac distances) similar to those of their inocula (the corresponding human stool sample), while the third sample was more different from the final SICs (Fig. 7D) due largely to the emergence of the Eubacteriaceae and Enterococcaceae from undetectable to 0.29±0.13% and 4.9±1.1%, respectively. Nonetheless, the final SICs derived from human stool after 7 passages were as (or more) similar to their inocula than the humanized mouse-derived SICs were to their inocula (the corresponding mouse fecal sample) using phylogenetic distance metrics (Fig. 7C,D). Taken together, these data demonstrate that *in vitro* batch culturing of human stool samples is a generally applicable tool to build complex, stable, and reproducible SICs.

## Discussion

Here, we have shown that *in vitro* passaging of fecal samples is a powerful tool for generating stable, host-relevant SICs in a deterministic and reproducible manner (Fig. 1G,H,S2A). Remarkably, this reproducibility even extends to a highly perturbative antibiotic treatment that causes large-scale compositional changes (Fig. S7B,G). SICs rapidly converge (Fig. 1D,S1L) and can be frozen (Fig. 1J) and cultured in larger volumes (Fig. 1I) without disrupting steady-state composition, demonstrating their utility for large-scale manufacturing and experimentation. Particular media may be useful for enhancing the abundance of certain desired families, while antibiotics and bacteriophages may be useful to remove undesirable taxa. Importantly, our observation that different inocula yield different SICs (Fig. 2, 7) showcases the potential of SICs for personalized therapeutics.

While our approach yields SICs that are more complex than ones typically studied with bottom-up approaches, the scaling of richness with growth yield (Fig. 2E) suggests that niche occupancy in BHI is saturated at ∼50 species, leading to competitive exclusion (Del Monte-Luna et al., 2004). It will be intriguing to investigate whether it is possible to achieve higher richness in other media, or by supplementing BHI with vitamins, cofactors, or specific nutrients such as mucin to promote the growth of fastidious organisms (Tramontano et al., 2018), like the Verrucomicrobia. Cross-feeding may increase richness and reshape community composition via niche generation (Goldford et al., 2018). Conversely, media composed of monomeric and dimeric carbon sources that are absorbed in the small intestine select for a few dominant species (Fig. S1K). A better understanding of the contributions of interspecies interactions and nutrient availability to community diversity and yield will be instrumental to engineering culturing systems to achieve the bacterial density and diversity observed in the gut. Our SICs represent a major step toward this goal.

Strikingly, while YCFA supports the growth of a wide range of gut commensals in pure cultures, in SICs the Enterococcaceae tend to take over (Fig. S1J, S3A), indicating that media that support the growth of many isolates do not necessarily support diverse SICs. Further, the ability of species to grow in isolation is generally not a good predictor of their relative abundance within an SIC (Fig. 3L). While the maximal growth rate of SICs scales with the fraction of fast growers (Fig. 2F), fast growers are quickly limited (Fig. 3M). Thus, fast growers probably dictate SIC growth rate, but final composition is determined by many other factors, which explains why BHI SICs are far more diverse than a scenario in which the rapid growers take over. Measurements beyond SIC composition—such as consumption and production of metabolites, growth and death rates of species, and global environmental properties such as pH and osmolarity—will enable a mechanistic understanding of community niche dynamics. Since SIC composition can be approximately mimicked with reference strains (Fig. I,K), incorporating genetically modifiable organisms and existing mutant libraries in our SICs could open additional applications.

A remarkable aspect of our SICs is their ability to recapitulate *in vivo* behaviors. Community composition was generally maintained through passaging in BHI (Fig. 1,7) and the microbial metaproteomic signature was similar between feces and SICs (Fig. S1E), suggesting that many interspecies interactions are conserved in this medium. Moreover, SICs derived from mice during and after antibiotic treatment were more susceptible to invasion by *S*. Typhimurium than SICs derived before treatment (Fig. 6A-C). While previous studies reported that ciprofloxacin treatment can increase the colonization susceptibility of chemostat-grown microbial communities (Carman et al., 2004), our experiments were conducted in the absence of ciprofloxacin at the time of challenge. Thus, the compositional changes elicited by antibiotics *in vivo* are maintained *in vitro*, leading to different functional outcomes.

Importantly, our study also allowed a quantitative comparison between the antibiotic response of the gut microbiota *in vivo* and *in vitro* (Fig. 5), which revealed remarkable similarities in taxon-specific changes in diversity (Fig. 5H). Many aspects of the recovery dynamics that are confounded by multiple factors *in vivo* were clarified by our *in vitro* studies. The lack of substantial recovery in richness *in vitro* after ciprofloxacin treatment (Fig. 5D) suggests that the ∼4-fold increase *in vivo* post-treatment relative to residual samples (Fig. S1B) comes from environmental reseeding or increased densities *in vivo*. Notably, SICs derived from feces at the peak or during residual treatment were more similar to their inocula than to feces post-treatment (Fig. 2A), indicating that removal of the drug in inoculating media *in vitro* is not enough to recover taxa that went undetectable *in vivo*. Highlighting the physiological relevance of such reservoirs, our *in vivo* studies indicate that singly housed mice experience a prolonged decrease in viable load, *Bacteroides* abundance, and diversity compared with co-housed mice during antibiotic treatment, with highly heterogeneous recovery dynamics across mice (Ng et al., 2019). Disruption of community composition and the set of species that become undetectable are mostly deterministic (Fig. S9C,H). Our findings show that the recovery of some species such as *B. fragilis* from undetectable levels across multiple passages does not require environmental reservoirs (e.g. via coprophagy in mice) or larger population sizes *in vivo*, indicating that resilience is built-in to some species, and pointing to SICs enabling the elucidation of aspects of community ecology that can be uncoupled from host dependence. Future work with fecal-derived SICs, including comparisons of our methodology with other fecal-derived and bottom-up reconstitution approaches (Macfarlane et al., 1998; Minekus et al., 1999; Van de Wiele et al., 2015), should elucidate the interplay among antibiotic sensitivity and tolerance, duration, and strength of antibiotic exposure, and migration between hosts and the environment.

In general, top-down and bottom-up SICs serve as powerful, complementary resources for the microbiota field. Bottom-up SICs can be used to measure interspecies interactions using drop-out experiments (Gutierrez and Garrido, 2019), while top-down SICs provide a more complex template for evaluating the effects of perturbations and comparing to *in vivo* results, including the potential for blooming of species from below the limit of detection pre-treatment. The overrepresentation of fast-growing species in SICs (Fig. 1) and their receding in SIC-colonized mice (Fig. S1C,D) shed light on the bacterial taxa whose growth is controlled by the host. Further, we predicted the effects of ciprofloxacin (Fig. 4G) at the ASV level through isolate-sensitivity measurements and perturbations to pre-treatment SICs. The emergence of resistant strains, and their out-competition by close relatives, can also be predicted (Fig. 6). The observed correlation between changes upon ciprofloxacin treatment and strain sensitivities (Fig. 4F) suggests that interspecies interactions are not a strong modifier of antibiotic responses in these SICs. Nonetheless, interspecies interactions can change antibiotic sensitivity in auxotrophic food-webs (Adamowicz et al., 2018) and induce tolerance (Aranda-Díaz et al., 2019). Notably, we can only measure isolate sensitivities in the culturable portion of the SICs (Fig. 3B), and metabolic dependencies of the strains we did not isolate could affect their survival. Further work will be needed to investigate the role of tolerance in the ability of specific taxa to recover after treatment (Fig. 5F,K). These findings suggest that first dissecting *in vitro* how a gut community is remodeled by a perturbation can serve as a powerful pre-screening tool before expensive mouse experiments.

Many other fascinating questions should be tackled in the future using our *in vitro* approach, which can be scaled to thousands of SICs simultaneously. Since a fecal sample from a mammalian host likely represents an average over the spatial localization and physiologies determined by the history and environmental contexts within the gut, a similar culturomics approach should be applied to samples from other locations in the gastrointestinal tract. Our observation that BHI-passaged SICs can shift the host proteome of germ-free mice toward that of humanized mice (Fig. S1D) suggests that SICs can also be safely re-introduced to their original hosts. Fecal microbiota transplantation is a promising therapy for diseases like *Clostridium difficile* infection (Drekonja et al., 2015) and aids in recovery after chemotherapy (Le Bastard et al., 2018), diarrhea (Tropini et al., 2018), and antibiotic treatments (Ng et al., 2019). However, the lack of standardization and danger due to potential collateral infections are major barriers for the wide therapeutic use of fecal microbiota transplantation (Cammarota et al., 2017). SICs may lower the risk of infection and improve manufacturing efficiency (Khoruts and Weingarden, 2014), thus overcoming these barriers to therapy. More generally, SICs should bridge mechanistic studies of microbial ecology and human-relevant *in vivo* systems to accelerate the pace of discovery in the microbiota field.

## Author Contributions

A.A.-D, K.M.N., J.L.S., and K.C.H designed the research; A.A.-D., K.M.N., T.T., S.D., F.B.Y., I.R.R., T.C., S.H, K.V., C.G.G., and T.N. performed the research; A.A.-D., K.M.N., D.D., and C.G.G. analyzed the data; and A.A.-D, J.L.S., and K.C.H wrote the paper and all authors reviewed it before submission.

## Acknowledgments

The authors thank Rebecca Culver, Carolina Tropini, Kali Pruss, Alice Cheng, Manohary Rajendram, Surya Tripathi, and Myles Bartholomew for technical assistance; and Rita Oliveira, Lisa Willis, Handuo Shi, and Po-Yi Ho for valuable comments on the manuscript. Some reference strains used in this study were kindly provided by Michael Fischbach and Denise Monack. A.A.-D. is a Howard Hughes Medical Institute International Student Research fellow, a Stanford Bio-X Bowes fellow, and a Siebel Scholar. The authors acknowledge funding from the Allen Discovery Center at Stanford on Systems Modeling of Infection (to K.M.N. and K.C.H.). J.L.S. and K.C.H. are Chan Zuckerberg Investigators.

## Methods

### Mouse sampling

All mouse experiments were conducted in accordance with the Administrative Panel on Laboratory Animal Care, Stanford University’s IACUC. Germ-free, female,Swiss-Webster mice were gavaged with a human fecal sample obtained from a healthy anonymous donor as previously described (Kashyap et al., 2013). The microbiota was allowed to equilibrate for 6-8 weeks before perturbation experiments commenced. Humanized mice were orally gavaged with antibiotics (Sigma Aldrich) dissolved in 200 μL water for 5 days. For experiments involving a dietary switch, mice were first fed a standard diet (Purina LabDiet 5010) rich in MACs, and then a defined low-MAC diet (Harlan TD.86489) in which the sole carbohydrates are sucrose (31% w/w), cornstarch (31% w/w), and cellulose (5% w/w). Mice were euthanized with CO_2,_ and death was confirmed via cervical dislocation.

Fecal pellets were sampled on ice and taken into an anaerobic chamber within 30 min. Human stool samples were collected from 3 healthy donors by scooping a small sample from a receptacle into a conical tube on ice. The samples were transferred to an anaerobic chamber within 5 min of sampling.

### SIC passaging, storage, and revival

Approximately 100 mg of stool samples from humanized mice or human donors were resuspended into 200 μL phosphate-buffered saline (PBS) in 1.5 mL microcentrifuge tubes. Samples were incubated in PBS for 10-15 min at room temperature to encourage large pieces of fecal matter to disintegrate. Samples were vortexed for 5 min and allowed to sit for 1 min to let food particles settle. One microliter of this resuspension was inoculated into 200 μL medium in 96-well polystyrene microplates (Greiner Bio-One). Each resuspension was inoculated into four media (Table S2) in triplicate. The plate was covered with optical film, with a small (∼0.5 mm) hole poked above each well, outside of the plate reader’s light path, to allow gas exchange.

Incubation and optical density (OD) measurements were performed with an Epoch 2 plate reader (BioTek) in an anaerobic chamber at 37 °C with continuous shaking and OD_600_ measured at 7.5-min intervals. Plates were incubated in the plate reader for 48 h and then 1 μL of this saturated culture was transferred into 200 μL of fresh medium in a new plate. We refer to a 48-h cycle from inoculation to saturation as one passage. Samples used in the passaging experiments in Figs. 1 and 2 were taken in consecutive days, hence the plate was removed from the plate reader every 24 h, to inoculate and/or passage corresponding SICs. On a given day, if the communities were not passaged, then the whole volume was transferred into a new plate.

After passaging, 50 μL of the cultures were mixed with 50 μL sterile 50% glycerol (v/v in water) in crimp vials, sealed, and stored at −80 °C for long-term storage. The remaining volume was stored in the 96-well plate at −80 °C with an aluminum seal until DNA was extracted. BHI SICs were revived for experimentation by inoculation of frozen glycerol stocks directly into 3 mL BHI. Revived communities were incubated at 37 °C for 48 h and passaged twice before experimentation by diluting 1:200 into fresh medium every 48 h.

### SIC colonization

Mice were humanized with a human fecal sample obtained from the same healthy anonymous donor as was used for the ciprofloxacin-treatment experiment (Fig. 1A). Four weeks after humanization, fecal pellets were collected from 11 mice housed across 3 cages. A mixture of the 11 fecal pellets was used to inoculate 3 mL BHI to derive an SIC as described above. The SIC was passaged every 48 h with a 1:200 dilution into fresh medium. After 5 passages, a frozen stock was made by mixing equal volumes of a saturated culture and 50% glycerol (v/v in water). The SIC was revived by inoculating the frozen stock into 3 mL BHI and passaging twice before using 200 μL of the saturated culture to gavage germ-free, female, Swiss-Webster mice. culture. Fecal pellets were sampled from the germ-free mice on the day that they were gavaged, on day 24 after colonization for proteome analysis, and on day 28 for 16S sequencing.

### Strains and growth media

Growth media are listed in Table S2. These media were chosen because of their ability to grow diverse microbes: the nutrient-rich Brain Heart Infusion (BHI) is widely used to culture fastidious microorganisms; Tryptone-Yeast extract-Glucose (TYG), which is often used to culture members of the *Bacteroides* genus that includes many prevalent human commensals; Gifu Anaerobic Medium (GAM; a mixture of animal, plant, and yeast digests and extracts supplemented with glucose and starch), which cultures individual species with relative yields similar to their abundances in human microbiotas (Rettedal et al., 2014); and Yeast extract, Casitone, and Fatty Acids (YCFA), which has been used to culture a wide range of human gut commensals (Browne et al., 2016). Isolated and reference strains are listed in Tables S1 and S3, respectively. All media and materials were made anaerobic by incubating in a custom anerobic chamber (Coy Labs) for 48 h before use.

### Growth analyses

Maximal growth rate was calculated as the maximal slope of ln(OD) with respect to time (calculated from a linear regression of a sliding window of 11 timepoints). The duration of lag phase was defined as the time at which a strain first reached its half-maximum growth rate.

### 16S rRNA sequencing

DNA was extracted from whole fecal pellets or 50 μL of culture with a DNeasy PowerSoil HTP 96 kit (Qiagen 12955-4). 16S rRNA amplicons were generated using Earth Microbiome Project-recommended 515F/806R primer pairs using the 5PRIME HotMasterMix (Quantabio 2200410) with the following program in a thermocycler: 94°C for 3 min, 35 cycles of 94 °C for 45 s, 50 °C for 60 s, and 72 °C for 90 s, followed by 72°C for 10 min. PCR products were cleaned, quantified, and pooled using the UltraClean 96 PCR Cleanup kit (Qiagen 12596-4) and Quant-iT dsDNA High Sensitivity Assay kit (Invitrogen Q33120). Samples were sequenced with 250- or 300-bp reads on a MiSeq (Illumina).

Samples were de-multiplexed with Qiime v. 1.9.1 (Caporaso et al., 2010) using the commands “split_libraries_fastq.py --rev_comp_mapping_barcodes -- rev_comp_barcode --store_demultiplexed_fastq --max_bad_run_length 999 -- min_per_read_length_fraction .01 --sequence_max_n 999 --phred_quality_threshold 0” and “split_sequence_file_on_sample_ids.py”. Subsequent processing was performed using DADA2 as previously described (Callahan et al., 2016). truncLenF and truncLenR parameters were set to 240 and 180, respectively. Resulting ASV sequences were assigned taxonomy with Qiime, using the commands “assign_taxonomy.py”, “align_seqs.py”, and “make_philogeny.py”. Custom MATLAB R2018a (MathWorks) scripts were used for further analysis.

ASVs were classified as contaminants and removed from further analysis if they appeared with significantly higher relative abundances (*p*<0.001, t-test) in control samples (water added to DNA extraction step).

Alpha diversity (richness) was measured by rarefying all samples to 5000 reads and calculating the number of taxa with relative abundance >0. Shannon diversity was calculated using 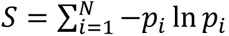, where *p* is the relative abundance of the *i*th taxon and *N* is the total number of taxa with *p*>0. Evenness was calculated using the Pielou index by normalizing Shannon diversity by the maximum Shannon index for the number of taxa: 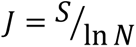.

Weighted and unweighted Unifrac distances between samples for beta diversity were calculated using custom MATLAB code. Unifrac was calculated as described in (Lozupone and Knight, 2005). Weighted Unifrac was calculated as described in (Lozupone et al., 2007): for samples *A* and *B*, 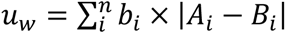, where *n* is the total number of branches in the tree, *b_i_* is the length of branch *i*, and *A_i_* and *B_i_* are the relative abundances of the taxa that descend from branch *i* in samples *A* and *B*, respectively. No further normalization was performed. Only taxa present at >0.01% in more than two samples were used to calculate distances. Principal coordinate analyses were performed on unweighted Unifrac distances between samples with >5000 reads.

### Isolation of strains

SICs were resuspended in PBS and plated onto BHI, BHI-S, or GAM plates. Plates were incubated at 37 °C in an anaerobic chamber. After 2 days, colonies were grown in the medium corresponding to the plate from which they were isolated for 2 days, and glycerol stocked. Strains were identified via high-throughput MALDI-TOF mass spectrometry of whole colonies, and subsequent matching of spectra to a reference library with a MALDI Biotyper System (Bruker), following manufacturer’s instructions and including formic acid lysis. For the SIC that we attempted to reconstitute, 15 distinct strains were obtained and their taxonomy was checked by Sanger sequencing. Genomic DNA was extracted from pure cultures using a DNeasy UltraClean 96 Microbial Kit (Qiagen 10196-4) or DNeasy Blood and Tissue Kit (Qiagen 69504) kit. The 16S gene was amplified using primers 5’AGAGTTTGATCCTGGCTCAG and 5’GACGGGCGGTGWGTRCA, and the amplicon was Sanger sequenced. Taxonomic assignment was performed by alignment using BLAST against the 16S ribosomal RNA sequences (Bacterial and Archaea) database.

### Community reconstitution

For community reconstitution, frozen stocks of isolated (Table S1) or reference (Table S3) strains were streaked onto BHI 5% sheep blood agar plates and incubated for 48 h at 37 °C in anaerobic conditions. A single colony of each strain was inoculated into 3 mL BHI in isolation, and incubated for 48 h at 37 °C. OD was measured for each strain and all were diluted to a final OD of 1×10^-4^ before mixing in 200 μL BHI in a 96-well plate. Mixed cultures were grown for 48 h in a plate reader to measure growth. After 48 h the cultures were diluted 1:200 into fresh BHI.

### Antibiotic sensitivity estimation

Frozen stocks of isolated strains (Table S1) were streaked onto BHI 5% sheep blood agar plates and incubated for 48 h at 37 °C in anaerobic conditions. A single colony of each strain was inoculated into a 3 mL monoculture in BHI, and incubated for 48 h at 37 °C. Cultures were then diluted 1:200 into fresh BHI containing 0.5-32 μg/mL ciprofloxacin. The minimum inhibitory concentration (MIC) for each strain was calculated as the minimum dose necessary to completely inhibit growth as measured by OD after 48 h of incubation.

### *In vitro* antibiotic treatment

SICs were revived in 3 mL BHI and passaged twice before experimentation by diluting 15 μL into 3 mL fresh medium every 48 h. One microliter of the SICs was then diluted into 200 μL fresh BHI containing 0.5-32 μg/mL ciprofloxacin. Growth was measured in a plate reader as described above, and 48 h-old cultures were stored at −80 °C until DNA was extracted.

### *In vitro S.* Typhimurium challenge

SICs were revived in 200 μL BHI in microplates and passaged twice by diluting 1 μL into 200 μL fresh medium every 48 h. A frozen stock of *S*. Typhimurium strain SL1344 was grown anaerobically at 37 °C for 24 h in BHI. One microliter of the SICs and 1 μL of *S*. Typhimurium adjusted to OD_600_=0.1 was diluted into 200 μL fresh BHI. Saturated co-cultures were serially diluted into sterile PBS after 48 h of incubation at 37 °C. Three microliters of a 1:10^4^ dilution were spotted onto LB agar with 50 μg/mL streptomycin to select for *S*. Typhimurium. Pictures of colonies (Fig. 6B, S8A) were taken after 48 h of incubation at 37 °C in aerobic conditions.

For single-cell quantification of *S*. Typhimurium levels, we followed the protocol described above to grow SICs and an *S*. typhimurium 14028s strain expressing cytoplasmic mCherry (Table S3) prior to co-culturing. SICs and *S*. Typhimurium were mixed at various ratios and incubated anaerobically for 48 h at 37 °C. Saturated cultures were removed from the anaerobic chamber and spotted onto PBS agar plates to perform high-throughput phase and fluorescence microscopy (Shi et al., 2017). Percentages of *S*. Typhimurium (Fig. 6C, S8B) were estimated by calculating the proportion of pixels within cell contours segmented from phase images whose fluorescence was above a threshold that clearly separated mCherry-positive and mCherry-negative cells using custom Matlab code.

### Isolation of stool proteins and peptides

Isolation of stool proteins broadly followed the protocol in (Gonzalez et al., 2020). Briefly, ∼50 mg mouse pellets or pelleted cells from 1 mL saturated cultures were aliquoted into a 96-well plate along with ∼600 mg of 0.1-mm ceramic beads (Omni International, #27-6006). To each well, 750 μL lysis buffer (6 M urea, 5% sodium dodecyl sulfate, and 50 mM Tris, pH 8.1) was added and plates were sealed with sealing mats (Omni International, #27-530). Sealed plates were subjected to 10 min of bead beating at 20 Hz using a Qiagen Tissuelyser II. After bead beating, each plate was centrifuged at 3000 rcf at 4 °C for 10 min. Five hundred microliters of the resulting supernatant were transferred to a new 2-mL 96-well plate (Waters, 186002482), sealed with a sealing mat, spun again at 3000 rcf at 4 °C for 10 min, then transferred to a fresh 2-mL plate. Samples were reduced with 10 μL of 500 mM dithiothreitol (Sigma-Aldrich) for 30 min at 47 °C and alkylated with 30 μL of 500 mM iodoacetamide (Sigma-Aldrich) for 1 h at room temperature in the dark. Fifty microliters of the reduced and alkylated supernatant were transferred to a new 2-mL 96-well plate for further processing, while the remaining material was stored at −80 °C.

Supernatant-resident stool proteins were washed, digested, and eluted as described in the Protifi S-trap protocol (http://www.protifi.com/wp-content/uploads/2018/08/S-Trap-96-well-plate-long-1.4.pdf). Briefly, the 50 μL supernatant was acidified with 5 μL of 12% phosphoric acid to which 300 μL S-trap binding buffer was added. Each resulting mixture was loaded into a single well. Positive pressure was used to load the proteins into each well with a Positive Pressure-96 Processor (Waters) with pressure set at 6-9 psi on the “Low-Flow” setting. Loaded proteins were washed with 300 μL Binding Buffer (90% methanol and 10% triethylammonium bicarbonate buffer (TEAB, Sigma-Aldrich, #T7408), adjusted to pH 7.1 using phosphoric acid) five times. After washing, 125 μL digestion buffer (100 mM TEAB and 5 μg trypsin) were added and proteins were digested for 3 h at 47 °C. Peptides were eluted with 100 μL TEAB, followed by 100 μL of 0.2% formic acid, followed by 100 μL of 50% acetonitrile, 0.2% formic acid. These peptides were captured in a 1-mL 96-well plate (Thermo Scientific, #AB-1127) and the volume was dried down in a Centrivap speedvac (Model 7810016). Plated samples were desalted using RP-S cartridges on an Agilent Bravo AssayMAP using the built-in desalting protocol, eluted with 50% acetontrile, and dried down. Plated peptide concentrations were normalized using a Nanodrop ND-1000.

### Mass spectrometry

Peptide samples were diluted to 0.5 μg/μL. Subsequently, 1 μL was loaded onto an in-house, laser-pulled 100-μm inner diameter nanospray column packed to ∼220 mm with 3-μm 2A C18 beads (Reprosil). Peptides were separated by reversed-phase chromatography on a Dionex Ultimate 3000 HPLC. Buffer A of the mobile phase contained 0.1% formic acid in HPLC-grade water, and buffer B contained 0.1% formic acid in acetonitrile. An initial 2-min isocratic gradient flowing 3% B was followed by a linear increase to 25% B for 115 min, then increased to 45% B over 15 min, and a final increase to 95% B over 15 min, whereupon B was held for 6 min and returned back to baseline in 2 min and held for 10 min, for a total of 183 min. The HPLC flow rate was 0.400 μL/min. Samples were run on a Thermo Fusion Lumos mass spectrometer that collected mass spectrometry data in positive-ion mode within the 400-1500 *m*/*z* range with an initial Orbitrap scan resolution of 120,000, followed by high-energy collision-induced dissociation and analysis in the orbitrap using “Top Speed” dynamic identification with dynamic exclusion enabled (repeat count of 1, exclusion duration of 90 s). The automatic gain control was set to 4e5 for FT full mass spectrometry and to 1e4 for ITMSn.

### Peptide/protein database searching

Mass spectra were searched using Proteome Discoverer 2.2 using the built-in SEQUEST search algorithm. Three FASTA databases were searched: Uniprot Swiss-Prot *Mus musculus* (taxon ID 10090, downloaded January 2017), a custom concatenated database containing proteome data for all 15 bacteria in the reconstituted SIC (Fig. 3B), and a database containing common preparatory contaminants. Target-decoy searching at both the peptide and protein level was employed with a strict false discovery-rate cutoff of using the Percolator algorithm built into Proteome Discoverer 2.2. Enzyme specificity was set to tryptic, with static peptide modifications set to cambamidomethylation (+57.0214 Da). Dynamic modifications were set to oxidation (+15.995 Da) and N-terminal protein acetylation (+42.011 Da). Only high-confidence proteins (*q* < 0.01) were used for analysis.

### Proteome analysis

Custom MATLAB R2018a (MathWorks) scripts were used to analyze protein abundances. *M. musculus* analyses were restricted to proteins that were 10 times more abundant in germ-free, humanized, or SIC-colonized mice samples than in any *in vitro* bacterial cultures in our dataset. Similarly, *B. fragilis* and *B. producta* proteome analyses were restricted to proteins that were 10 times more abundant in humanized mice, SIC-colonized mice, and SIC samples than in germ-free mice.

