## Supplementary Information for "High-throughput cultivation of stable, diverse, fecal-derived microbial communities to model the intestinal microbiota"

**Supplementary Text**

***SICs are reproducible across media and inocula***

We computed Pearson correlation coefficients of amplicon sequence variant (ASV) abundances among technical replicates for all combinations of inoculum and medium, and found that 90% were >0.68 (Fig. S1E), with a mean and standard deviation of 0.86±0.22 ($n$=192). Across all pairwise comparisons, the mean number of unique ASVs (>0.1% in one replicate and not detectable in the other) was very low (3.0±3.3, $n$=384), with a mean $\log_{10}$(relative abundance) of -2.24±0.62 ($n$=1171), suggesting that many instances of unique ASVs were due to abundances lower than the limit of detection rather than due to stochasticity in passaging. SICs cultured in YCFA had the largest proportion of replicates with low correlation coefficients (<0.6; Fig. S1E,F), while every pair of replicates cultured in BHI had a correlation coefficient >0.6 (Fig. S1F). SICs with negative correlation coefficients were cultured from antibiotic-treated inocula (SD mouse 2, residual and MD mouse 2, peak) in GAM or YCFA, and the negative correlation was explained by the dominance or absence of the Enterococcaceae family (Fig. S1G). Half of the technical replicates shared members that constituted >99% of the total abundance, while only ~1% had shared members accounting for <5%. In general, the lowest replicability was observed in YCFA and in SICs cultured from inocula with low diversity (Fig. S1G), suggesting that higher diversity promotes deterministic behavior during passaging. Intriguingly, these findings indicate that SICs are not generally stable in GAM and YCFA, despite the ability of these media to promote the growth of many commensals in isolation.

***SIC and fecal-sample richness are highly correlated due to maintenance of abundant species and within-family replacement***

Since BHI-passaged SICs were more compositionally similar to their inocula than SICs derived in other media (Fig. 2A), we focused on these SICs and queried the extent to which their richness was due to species that were detectable in the inoculum as opposed to emergent species that increased above the limit of detection due to passaging. We first focused on the three replicate SICs derived from the pre-treatment MD mouse sample in Fig. 1D-G. The fraction of ASVs detectable in the fecal sample that were detectable on average beyond the fourth passage was 39.1±2.6% (23/58 ASVs). The fractions of families and genera that were detectable after passaging were higher and similar to each other (52.9% and 56.0%, respectively), suggesting that loss of a genus often meant the loss of an entire family. Of the lost ASVs, their median $\log_{10}$(relative abundance) in the inoculum was lower than that of the retained ASVs (-2.46 and -2.05, respectively; $p$=0.006 Wilcoxon test) and they constituted 55.1% of the inoculum, of which 38.0% corresponded to *A. muciniphila*, which has been shown to be fastidious (Tramontano et al., 2018). Lost ASVs were lost early in the culturing experiment, in

1.6±0.7 passages. Of the 37 ASVs lost in at least one replicate, 92% were lost in all three replicates, suggesting that ASV loss was not due to bottleneck effects of dilution during passaging. Thus, even though family-level composition is preserved for families conducive to culturing in these conditions, passaging leads to the deterministic decrease– biased toward species that start at lower abundance – of species to levels below the limit of detection.

While almost 40% of the ASVs detected in the inoculum were maintained, there were also 22.0±3.0 ASVs detected in SICs that were undetectable in the inoculum, which could be due to contamination or to the promotion during passaging of a jump in abundance from below to above the limit of detection. The lack of contamination in the control wells in our plates, along with the observation that replicates exhibited many of the same emergent ASVs (63% of the 27 emergent ASVs were present in all three replicates), suggested that contamination was generally low. These observations are reminiscent of the emergence of *A. muciniphila* within mice switched from a standard to MAC-deficient diet, although these bacteria were often undetectable before the dietary switch (Earle et al., 2015). Most emergence in our SICs was due to within-family replacement; the only emergent family was the sole Gammaproteobacteria in the SIC (an Enterobacteriaceae) that had no detectable members in the inoculum. The ASVs that emerged typically did so in all three replicates, further underscoring the deterministic nature of SIC dynamics even at low abundance. As a whole, emergent ASVs accounted for 44.6±0.9% of the SIC, with 29.1±2.5% due to the Enterobacteriaceae ASV, and their median $log_{10}$(relative abundance) in the SICs was similar to that of the ASVs that were retained (-2.13 and -2.14, respectively; $p$=0.35 Wilcoxon test). Emergent ASVs reached stable levels, defined as within 2 standard deviations of their mean $log_{10}$(relative abundance) beyond the fourth passage, in 2.05±1.22 passages. These data suggest that the emergence of an ASV does not predict its abundance in the SIC and that restructuring of SICs (both loss and emergence) primarily occurs in the first two passages.

The total number of ASVs present in at least one inoculum (fecal gamma diversity) was 158, of which 68 were present in at least one BHI SIC and 79 in any medium. The total number of ASVs present in at least one SIC was 117. Of these, two appeared in at least one technical replicate of all BHI SICs, an *Enterococcus* species and a member of the Lachnospiraceae family; four more Lachnospiraceae were included when we ignored the low-diversity SIC cultured from residual treatment inocula. Intriguingly, these ASVs were present in the core microbiota of almost all mice at the point of minimum alpha diversity during antibiotic treatment (Ng et al., 2019), suggesting their general ability to persist through perturbations.

Species loss and emergence were ubiquitous across all SICs in all media. The fraction of ASVs detectable in the fecal sample that remained detectable in the seventh passage was negatively correlated with the diversity of the inoculum (for BHI, $R$=-0.89, $p$=$10^{-17}$; Fig. S4B), consistent with our observation that less-abundant members are more likely to disappear. Conversely, the fraction of emergent ASVs was positively correlated with the diversity of the inoculum ($R$=-0.64, $p$=$10^{-6}$, Fig. S4C). However, the fraction of families lost was only weakly correlated with inoculum diversity (Fig. S4D), and the fraction of families that emerged was weakly anticorrelated with inoculum diversity (Fig. S4E), indicating general species replacement within families. Almost 90% of the ASVs that were lost or gained did so within the first four passages (Fig. S4F,G). Lost ASVs were present in as many as 14 of the 16 inocula, with a mean of 3.9±3.4 inocula, and >66% went undetectable from all the inocula in which they were present, indicating that most of the disappearance dynamics are deterministic. While *A. muciniphila* was present in all inocula, it was maintained in 16.7% of all SICs and in 31.3% of BHI SICs, albeit at decreased abundance relative to the inoculum.

### Supplementary Figures

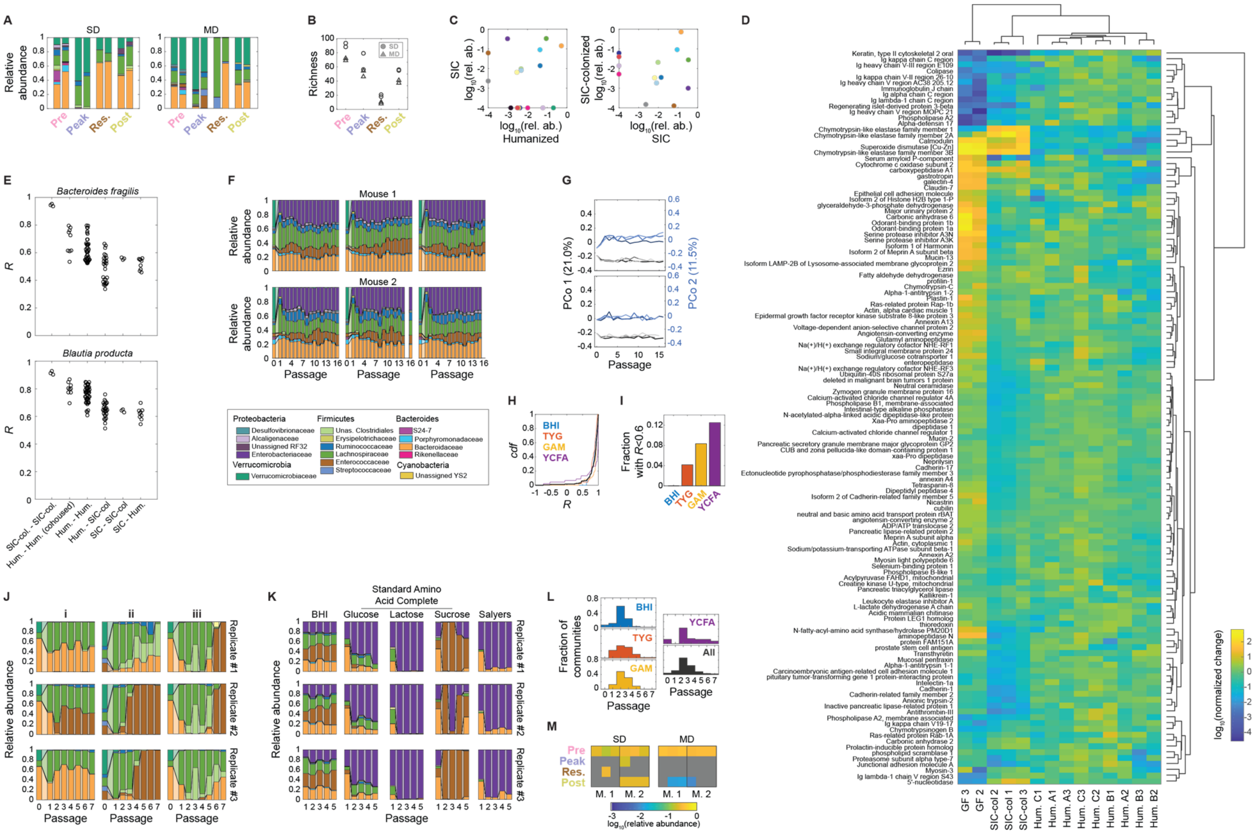


**Figure S1: Robust, high-throughput cultivation of fecal-derived SICs. Related to Figure 1.**

A) Ciprofloxacin elicits large changes in microbiota composition *in vivo*. Family-level composition of fecal samples from mice in Fig. 1A,B. Each timepoint (described in Fig. 1A) has two bars corresponding to the two mice in each group. Legend is located underneath (F). SD, standard diet; MD, microbiota-accessible carbohydrate-deficient diet.

B) Diversity *in vivo* decreases during ciprofloxacin treatment and does not fully recover after treatment. Richness (number of ASVs in rarefied data) of fecal inocula.

C) Families that are overrepresented *in vitro* recede *in vivo*. Shown is a comparison of family-level $log_{10}$(relative abundance) of an SIC and family-level $log_{10}$(relative abundance) of the humanized-mouse fecal inoculum from which it was derived (left), or ex-germ-free mice colonized with the SIC (right, SIC *n*=1, inoculum *n*=1, SIC-colonized *n*=3). Families with relative abundance of 0 were set to $10^{-4}$ for visualization. Mouse data are the same as in Fig. 1F.

D) The secreted proteome of mice colonized with SIC is similar to the proteome of humanized mice. *M. musculus* proteins present 10-fold higher in mice than in SICs were normalized by their mean abundance in humanized mice. Dendrograms resulting from hierarchical clustering of normalized relative abundance of proteins in germ free (GF), SIC-colonized (SIC col.), or humanized mice (Hum.) housed in three cages (A, B, and C).

E) Metaproteome of abundant species is similar *in vitro* and *in vivo*. *B. fragilis* or *B. producta* proteins present 10-fold higher in SICs than in germ-free mice were used to calculate the Pearson correlation coefficient of $log_{10}$(relative abundances) between samples.

F) *In vitro* passaging leads to stable and complex SICs. Family-level composition of three replicate SICs from two fecal samples (pre-treatment MD mice #1 and #2) during passaging in BHI for 16 rounds *in vitro*. Passage 0 is the fecal inoculum.

G) SICs rapidly converge to stable states. First (black) and second (blue) principal coordinates (PCo) of SICs in (C) over passages. The three replicates are depicted in different shades of the same color.

H) Technical replicates are largely reproducible. Cumulative density function of Pearson correlation coefficient (*R*) for all pairwise comparisons between technical replicates.

I) Proportion of technical replicate pairwise correlations $R<0.6$ for the 4 growth media.

J) Non-reproducible technical replicates share similar dynamics during early passages. Family-level composition during *in vitro* passaging for 7 rounds of three SICs with low correlation coefficients after 7 passages. (i) Technical replicates of SIC originating from SD mouse #2 during pre-treatment, grown in GAM. (ii) Technical replicates of SIC originating from MD mouse #2 during peak of treatment, grown in YCFA. (iii) Technical replicates of SIC originating from SD mouse #2 during residual treatment, grown in YCFA.

K) SICs derived in defined media are dominated by Enterococcaceae or Enterobacteriaceae. Family-level composition during 5 rounds of *in vitro* passaging for SICs originating from SD mouse #2 during pre-treatment, grown in Standard

Amino Acid Complete media supplemented with glucose, lactose, or sucrose, or in Salyers media supplemented with glucose.

L) The composition of an SIC generally converges in the first 4 passages. Histogram of passages required for SIC convergence (defined as weighted Unifrac distance between two consecutive passages <0.15).

M) While undetectable in the inocula, Enterobacteriaceae were found in all BHI-grown SICs derived from pre-treatment inocula but not in all SICs derived from inocula during and after treatment. Heatmap of $log_{10}$(relative abundance) of the Enterobacteriaceae in the seventh passage of SICs grown in BHI.

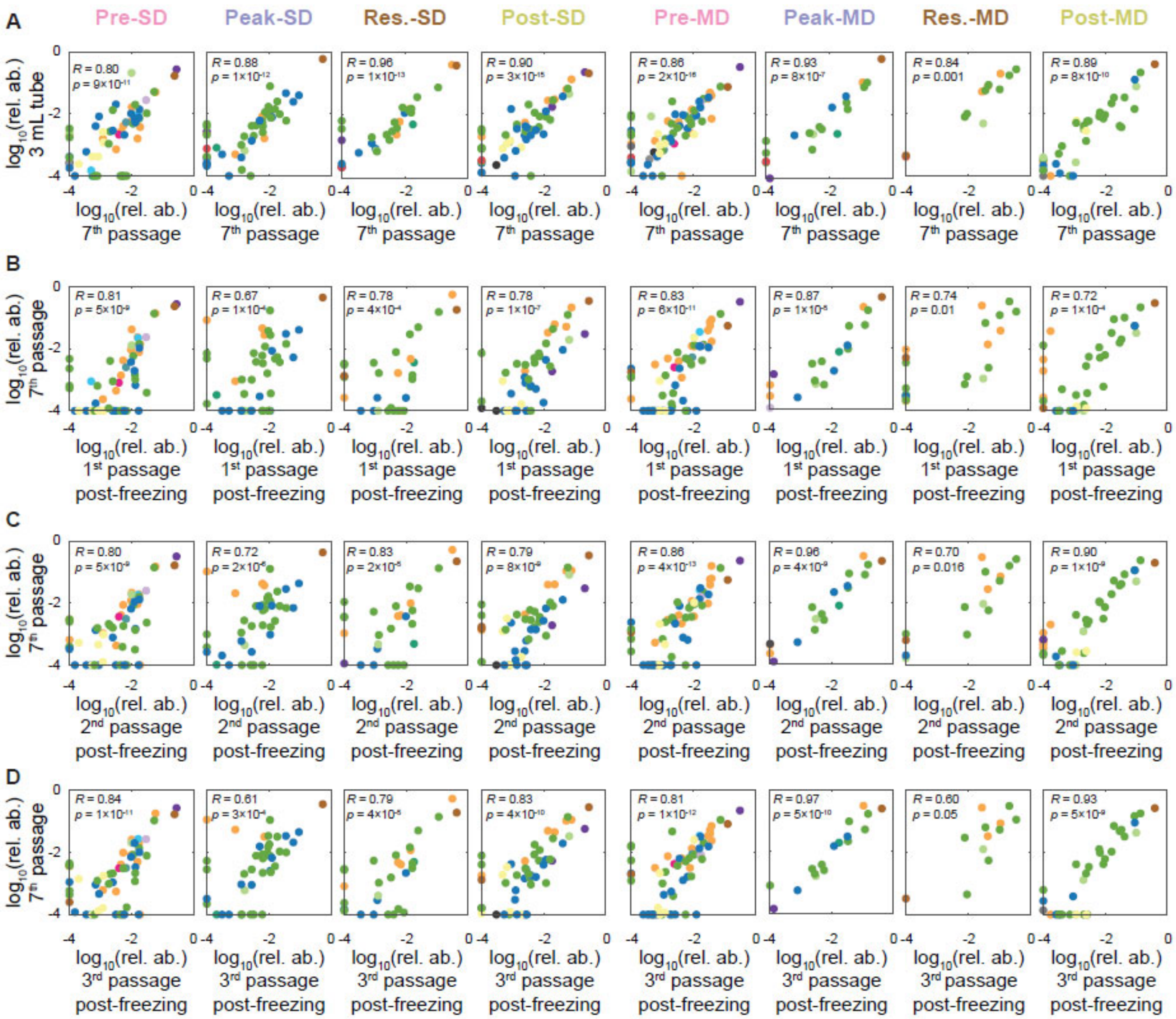


**Figure S2: SICs maintain composition after growth in larger volumes without shaking and during revival after freezing. Related to Figure 1.**

A-D) Correlation plots of $\log_{10}$(relative abundance) at the ASV level for 8 SICs grown in BHI after 7 passages against the same SIC grown for one passage in a larger volume (3 mL) without shaking (A), and after freezing and reviving for one (B), two (C), or three (D) passages. Pearson coefficients (*R*) and their *p*-values were computed using only data points present in both samples. ASVs with relative abundance of 0 were set to $10^{-4}$ for visualization purposes.

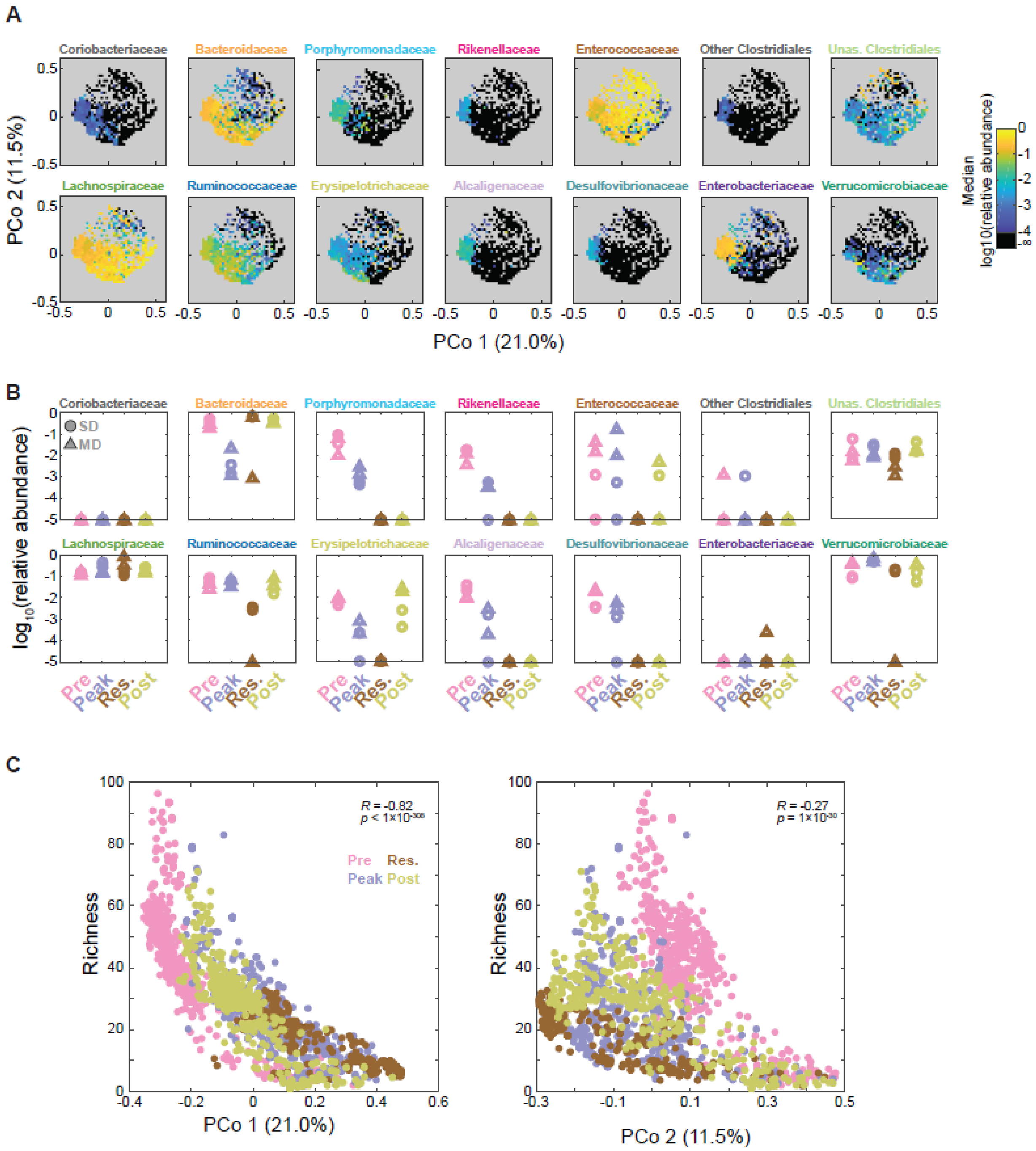


**Figure S3: Certain families generally co-occur in SICs, and SIC richness is strongly correlated with the first principal coordinate. Related to Figure 2.**

A) Median $\log_{10}$(relative abundance) of families across all samples (*n*=1728) binned by first two principal components. Only families present in >10% of the samples are shown.

B) $\text{Log}_{10}$(relative abundance) of families in (A) in each of the inocula. Families with relative abundance of 0 were set to $10^{-4}$ for visualization purposes.

C) Sample diversity was negatively correlated with each of the first two principal coordinates, particularly PCo 1, regardless of initial inoculum. Richness (number of ASVs in rarefied data) is plotted against the value of the first two principal coordinates for each of the samples (*n*=1728). Pearson correlation coefficients (*R*) and their *p*-values are reported.

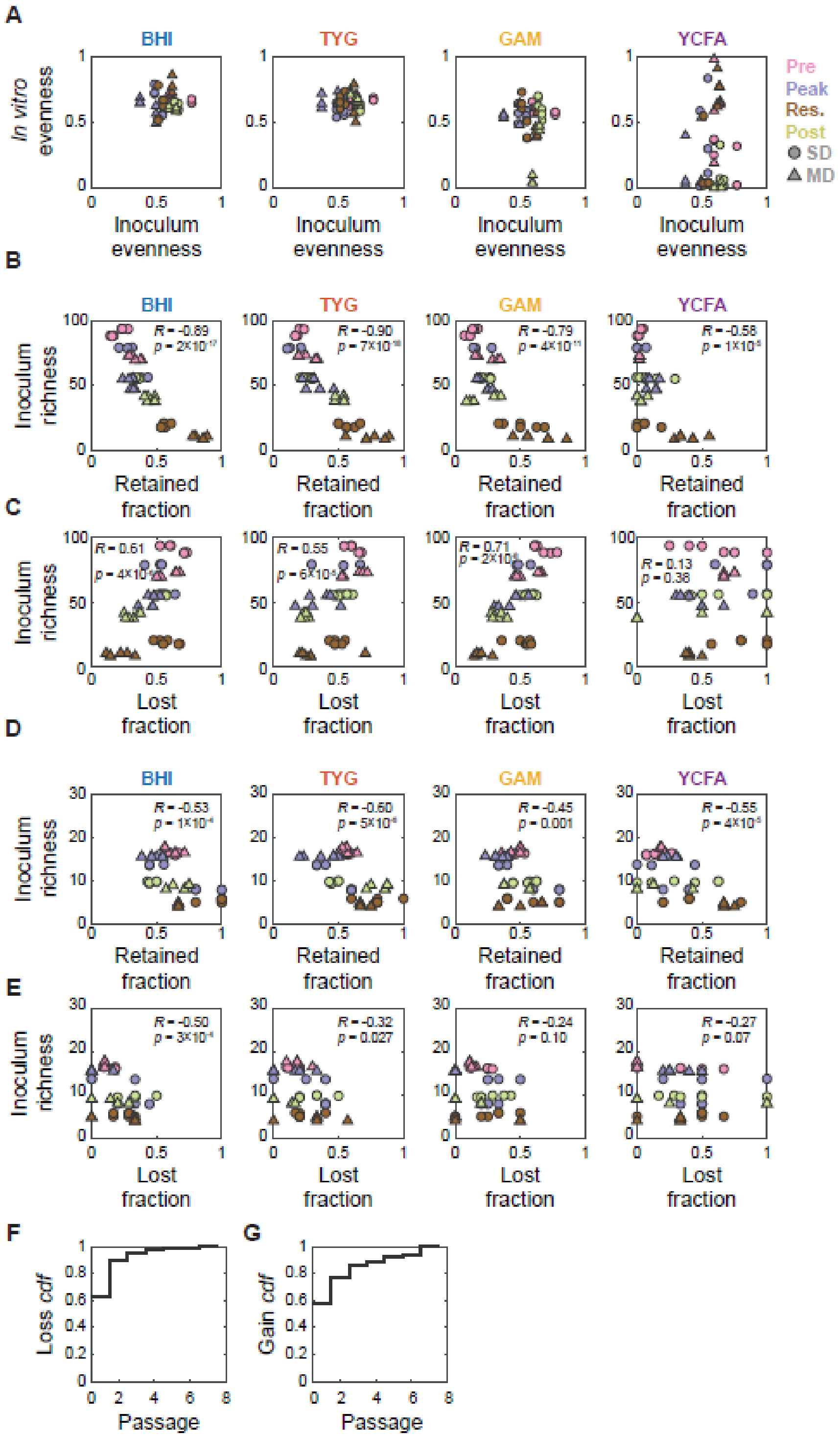


**Figure S4: Taxa loss and emergence are correlated with inoculum diversity. Related to Figure 2.**

A) Most growth media harbored SICs with evenness similar to that of the fecal inoculum from which each SIC originated. Evenness (Pielou's evenness index or normalized diversity) of the passaged SICs against inoculum evenness.

B-E) Correlation plots for inoculum diversity (number of ASVs in (B) and (C), and families in (D) and (E), in rarefied data) with fraction of taxa lost in (B) and (D) and emerged in (C) and (E), for all 192 passaged SICs. Pearson correlation coefficients (*R*) and their *p*-values were computed from all data points in each plot ($n$=48).

F-G) Most ASV loss or emergence occurred within the first three passages. Cumulative density function for the passage at which an ASV was lost (F) or emerged (G).

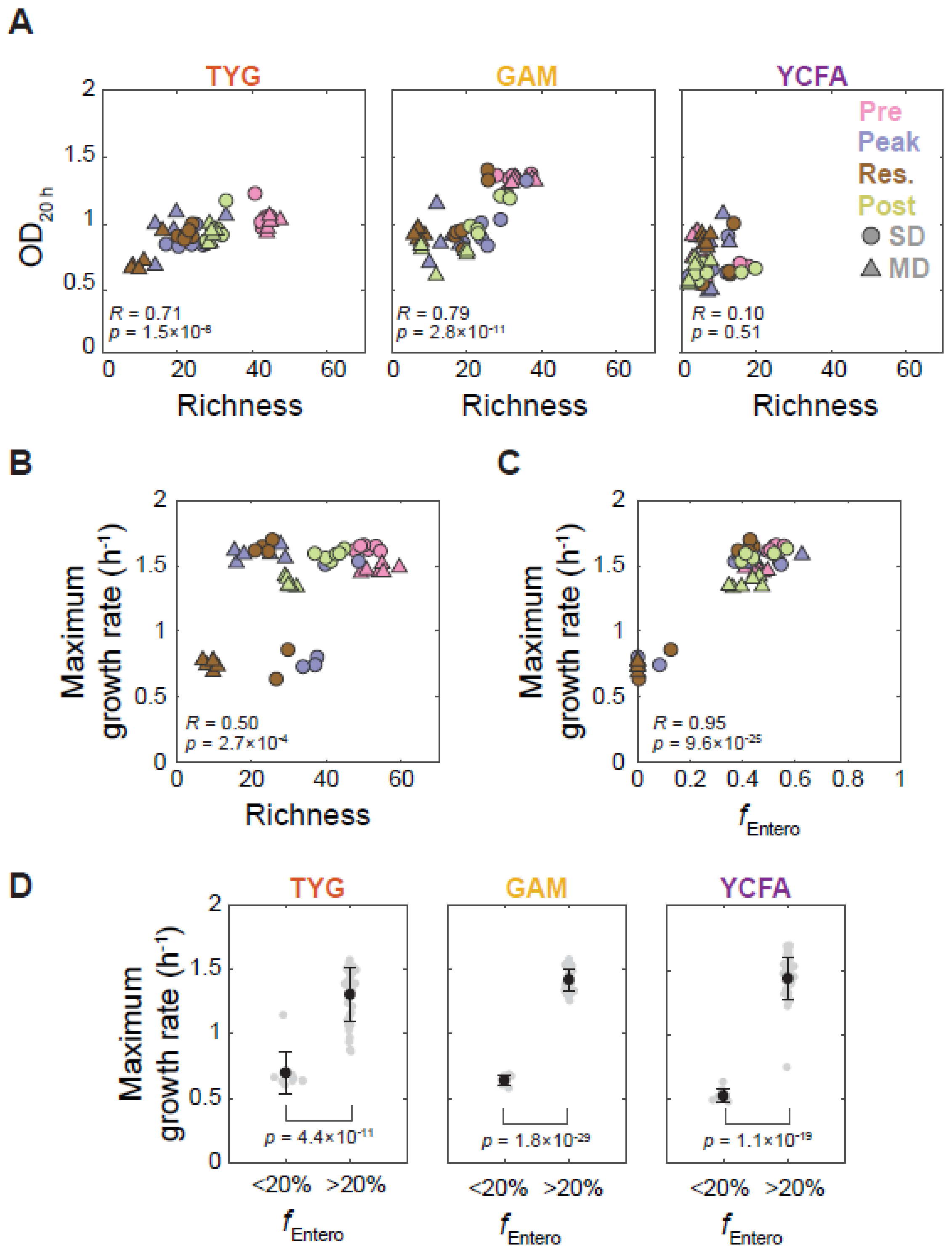


**Figure S5: SIC growth is linked to composition. Related to Figure 2.**

A) SIC yield scales with SIC diversity in most media. Optical density (OD) after 20 h of growth against richness (number of ASVs in rarefied data) of SICs grown in TYG, GAM, or YCFA. $OD_{20\ h}$ values are the mean of passages 3 to 7. Richness corresponds to the community in the seventh passage. Data points are colored and shaped corresponding to the fecal sample from which they originated. Pearson correlation coefficients ($R$) and their $p$-values are reported, $n$=48.

B) SIC maximum growth rate scales with SIC diversity. Maximum growth rate in seventh passage against richness (number of ASVs in rarefied data) of SICs grown in BHI. Pearson correlation coefficient ($R$) and its $p$-value are reported ($n$=48).

C) SIC maximum growth rate in BHI correlates with the presence of quickly growing families. Maximum growth rate in the seventh passage against the summed relative abundance of Enterobacteriaceae and Enterococcaceae ($f_{\text{Entero}}$). Data points are colored and shaped as in (B). Pearson correlation coefficient ($R$) and its $p$-value are reported ($n$=48).

D) Maximum growth rate is correlated with presence of quickly growing families in all media. Maximum growth rate was calculated from growth curves of the seventh passage and SICs were classified based on the summed relative abundance of Enterobacteriaceae and Enterococcaceae ($f_{\text{Entero}}$) for all SICs grown in TYG, GAM, or YCFA. Black circles are the mean maximum growth rate for each group (TYG: $n$=10 for $f_{Entero} < 20\%$ and $n$=38 for $f_{Entero} > 20\%$; GAM: $n$=9 for $f_{Entero} < 20\%$ and $n$=39 for $f_{Entero} > 20\%$; YCFA: $n$=8 for $f_{Entero} < 20\%$ and $n$=40 for $f_{Entero} > 20\%$), error bars represent standard deviation. Individual data points are plotted in grey. $p$-value is from a Student's two-sided $t$-test between the two groups.

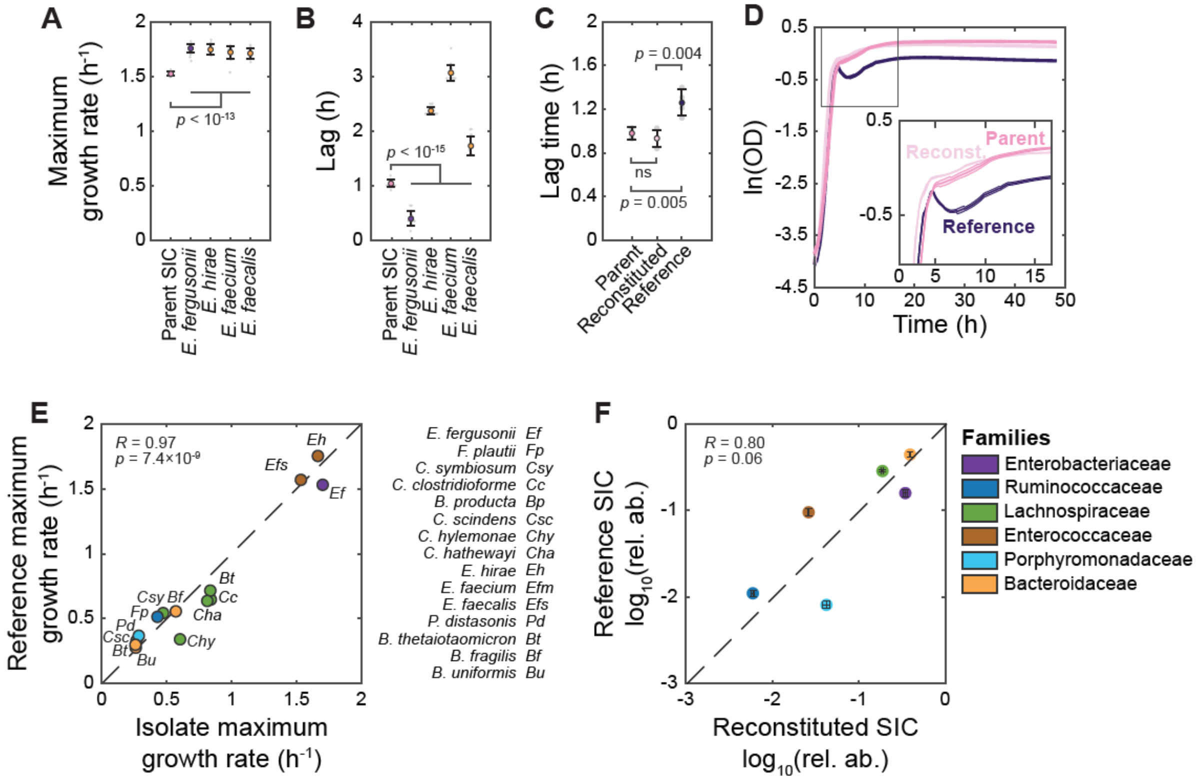


**Figure S6: Growth characteristics of SIC are distinct from those of its constituents. Related to Figure 3.**

A,B) Maximum growth rate (A) and lag time (B) of the parent SIC in Fig. 3, by comparison with the four fastest-growing isolated strains. Colored circles and error bars represent mean values and standard deviations (SD), respectively ($n$=4). $p$-values are from a Student's two-sided $t$-test between each pairwise comparison.

C) SIC growth characteristics are partially recapitulated by reconstituted and reference SICs. Lag time duration of the parent, reconstituted, and reference SICs. Symbols are as in (B); Colored circles and error bars represent mean values and SD, respectively (*n*=4). *p*-values are from a Student's two-sided *t*-test between each pairwise comparison; ns: not significant.

D) Differences in growth between SICs emerge upon entry to stationary phase. Inset: zoom-in to boxed region. Thick lines and thin lines above and below them represent the mean values and SD, respectively (*n*=4).

E) Maximum growth rates of isolates in Fig. 3C-E are highly correlated with those of the corresponding reference strains. *R* is the Pearson correlation coefficient (*n*=15).

F) Reconstituted and reference SICs converge to compositions with similar family-level relative abundances. Error bars are SD (*n*=4). *R* is the Pearson correlation coefficient (*n*=6).

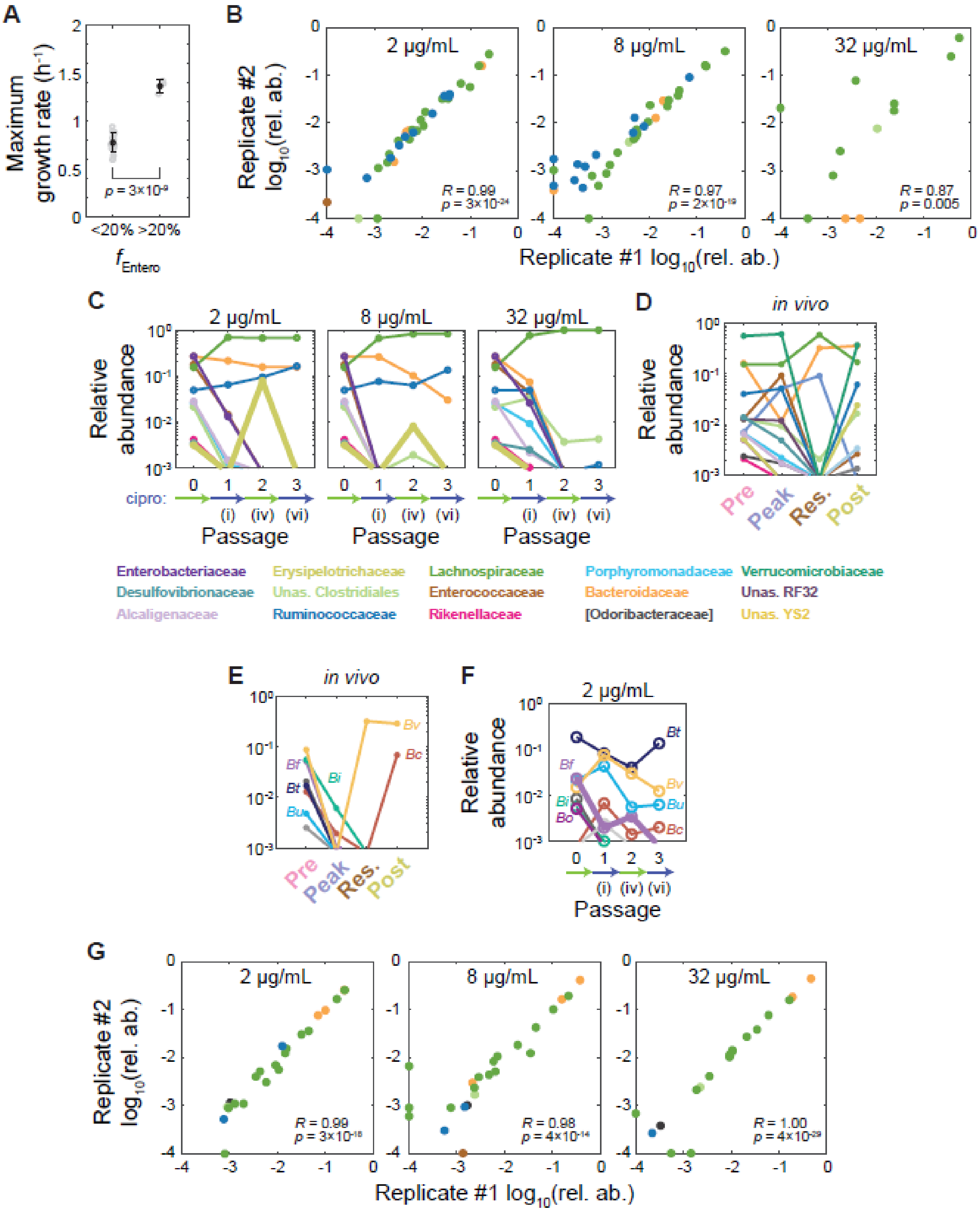


**Figure S7: Ciprofloxacin treatment restructures SICs. Related to Figures 5 and 6.**

A) Growth rate recovery after transient ciprofloxacin treatment is linked to the recovery of fast-growing species. Maximum growth rate of SICs grown in BHI after one round of treatment with ciprofloxacin and two rounds without drug. SICs were classified by their summed relative abundances of Enterococcaceae and Enterobacteriaceae ($f_{Entero}$). Black circles are the mean maximum growth rate for each group ($n$=21 for $f_{Entero} < 20\%$ and $n$=3 for $f_{Entero} > 20\%$), error bars are standard deviations (SD). Individual data points are plotted in gray. $p$-value is from a Student's two-sided $t$-test between the two groups.

B) Treatment outcome of Pre-SD SICs is reproducible, especially at low concentrations. Shown are comparisons of $\log_{10}$(relative abundance) at the ASV level between replicates after 3 passages of growth in BHI with ciprofloxacin. Pearson coefficient ($R$) and its $p$-value were computed only from data points present in both samples. ASVs with relative abundance of 0 were set to $10^{-4}$ for visualization purposes.

C) Erysipelotrichaceae recovery is reversed by a second ciprofloxacin treatment. Data are the family-level mean $\log_{10}$(relative abundance) of two replicates during one round of ciprofloxacin treatment followed by one round of recovery and a second treatment.

D) Ciprofloxacin-induced changes to family-level abundances in MD mice *in vivo* resemble changes in the corresponding SIC. Data are the mean $\log_{10}$(relative abundance) of two mice.

E) *Bacteroides* dynamics in MD mice consist of *B. vulgatus* dominance during treatment and the recovery of *B. caccae* after treatment. Data are the mean $\log_{10}$(relative abundance) of two MD mice at the ASV level for *Bacteroides* strains. *B. vulgatus* (*Bv*), *B. uniformis* (*Bu*), *B. caccae* (*Bc*), *B. intestinalis* (*Bi*), *B. fragilis* (*Bf*), and *B. thetaiotaomicron* (*Bt*). Other *Bacteroides* are shown in shades of grey.

F) *B. fragilis* recovery was reverted by a second ciprofloxacin treatment. Data are the ASV-level mean $\log_{10}$(relative abundance) of two replicates during one round of ciprofloxacin treatment followed by one round of recovery and a second treatment. *B. ovatus* (*Bo*), *B. vulgatus* (*Bv*), *B. uniformis* (*Bu*), *B. caccae* (*Bc*), *B. intestinalis* (*Bi*), *B. fragilis* (*Bf*), and *B. thetaiotaomicron* (*Bt*). Other *Bacteroides* are shown in shades of grey.

G) Treatment of the residual treatment humanized mouse fecal inoculum (Res-SD) SIC led to highly reproducible outcomes. Shown are comparisons of $\log_{10}$(relative abundance) at the ASV level between replicates after 3 passages of growth in BHI with ciprofloxacin. Pearson coefficient (*R*) and its *p*-value were computed only from data points present in both samples. ASVs with relative abundance of 0 were set to $10^{-4}$ for visualization.

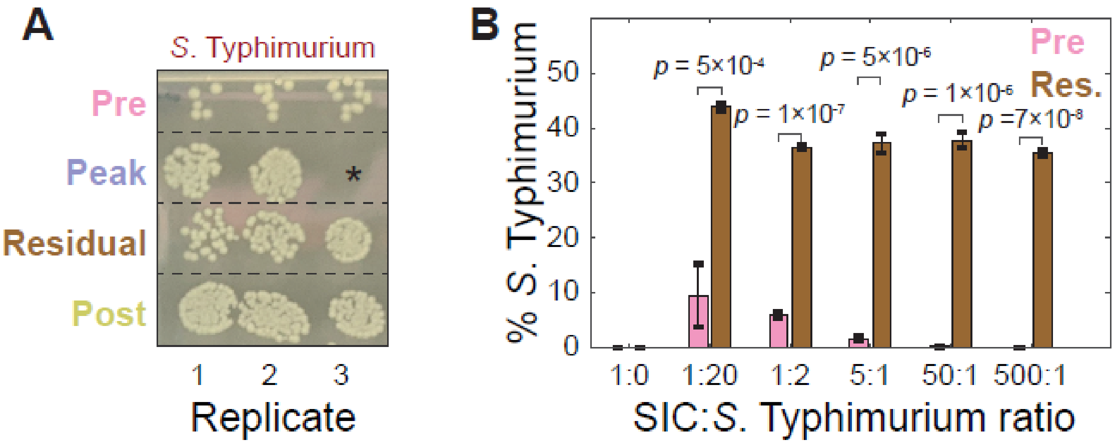


**Figure S8:** ***In vivo*** **ciprofloxacin treatment makes SICs more susceptible to** ***S.*** **Typhimurium invasion. Related to Figure 6.**

A) Colonies of *S*. Typhimurium SL1344 after 48 h of growth with SICs spotted on LB+streptomycin in aerobic conditions after a 1:$10^4$ dilution. These data are a biological replicate (SICs derived from mice housed in a different cage) of Fig. 6B. *missing replicate.

B) Single-cell quantification of mCherry-tagged *S*. Typhimurium 14028s after 48 h of co-culturing with SICs derived from pre- and residual-treatment mice fecal inocula. *p*-values are from a Student's two-sided *t*-test between each pairwise comparison, $n$=3.

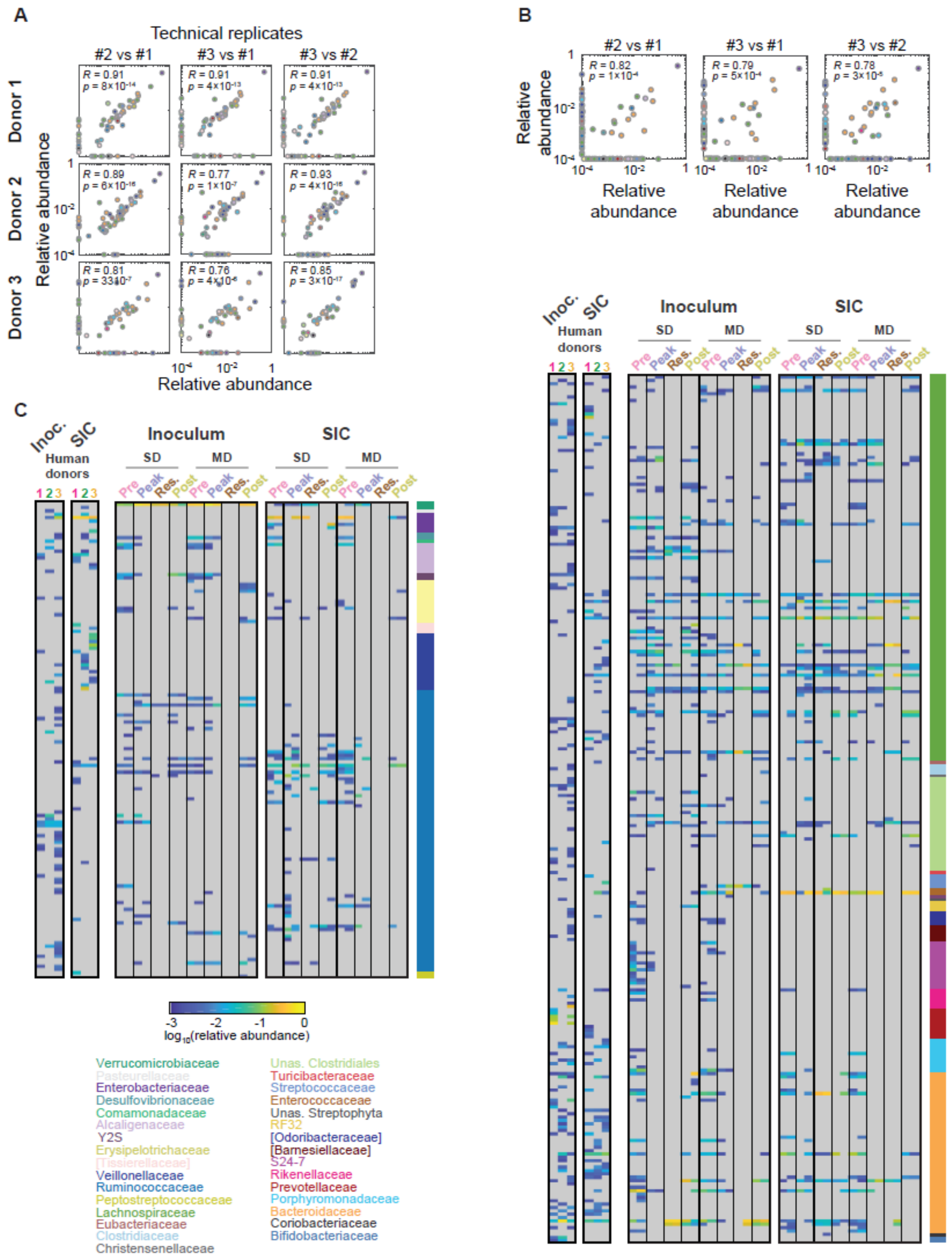


**Figure S9: SICs derived from human stool samples were highly reproducible, and ASVs in common overlapped with other SICs. Related to Figure 7.**

A) *In vitro* SICs derived directly from human stool were highly reproducible. Shown are comparisons of ASV-level $\log_{10}$(relative abundance) for three technical replicates of the SIC in Fig. 7A after 7 passages. ASVs with relative abundance of 0 were set to $10^{-4}$ for visualization. *R* and *p* were computed using only ASVs present in both samples.

B) SICs derived from different donors were distinct (Fig. 7A), but the abundances of ASVs in common between pairs of donors were highly correlated. Shown are comparisons of ASV-level mean $\log_{10}$(relative abundance) across three technical replicates of the SICs derived from each of the 3 human donors after 7 passages. ASVs with relative abundance of 0 were set to $10^{-4}$ for visualization. *R* and *p* were computed using only ASVs present in both samples.

C) *In vitro* passaging of human donor and humanized mouse fecal samples led to a collection of distinct SICs that maintained many families from the inocula. Shown are mean $\log_{10}$(relative abundance) across three technical replicates for each inoculum and the corresponding SIC after 7 passages. Only ASVs with mean relative abundance >0.1% are shown; ASVs with mean relative abundance <0.1% are shown in grey.

### Supplementary Tables

| **Strain** | **Closest relative in NCBI 16S ribosomal RNA sequence database** | **Ciprofloxacin MIC (μg/mL)** |
|---|---|---|
| TT1 | *Enterococcus hirae* ATCC 9790 | 2 |
| TT2 | *Escherichia fergusonii* ATCC 35469 | <0.5 |
| TT3 | *[Clostridium] symbiosum* ATCC 14940 | 16 |
| TT4 | *Bacteroides thetaiotaomicron* JCM 5827 | 16 |
| TT5 | *[Clostridium] clostridioforme* ATCC 25537 | 32 |
| TT6 | *Blautia producta* JCM 1471 | 32 |
| TT7 | *[Clostridium] scindens* strain DSM 5676 | 32 |
| TT8 | *Enterococcus faecium* strain DSM 20477 | 4 |
| TT9 | *[Clostridium] hylemonae* TN-272 | 32 |
| TT10 | *Enterococcus faecalis* NBRC 100480 | 4 |
| TT11 | *[Clostridium] hathewayi* 1313 | 32 |
| TT12 | *Bacteroides fragilis* NCTC 9343 | 4 |
| TT13 | *Flavonifractor plautii* 265 | 8 |
| TT14 | *Bacteroides uniformis* JCM 5828 | 16 |
| TT15 | *Parabacteroides distasonis* ATCC 8503 | 4 |

**Table S1: Strains isolated in this study.**

| **Medium** | **Formulation** | **Sterilization** | **Storage** |
|---|---|---|---|
| BHI | Commercially available (BD 211061) | Autoclave 20 min | Room temperature (RT) |
| TYG | As described in (Whitaker et al., 2017) | 0.22-μm filter | 4 °C |
| GAM | Commercially available (HiMedia M1801) | Autoclave 20 min | RT |
| YCFA | As described in (Duncan et al., 2002). Vitamins and cysteine added 48 h before experiment | 0.22-μm filter | 4 °C |

**Table S2: Media used in this study.**

| Strain | Closest isolate in this work | Source |
|---|---|---|
| *Enterococcus faecium* TX1330 | TT1 and TT8 | Huang lab |
| *Escherichia fergunsonii* DSM | TT2 | Michael Fischbach's lab, Stanford |
| *[Clostridium] symbiosum* WAL-14673 | TT3 | Huang lab |
| *Bacteroides thetaiotaomicron* VPI-5482 | TT4 | Sonnenburg lab |
| *[Clostridium] clostridioforme* 2_1_49FAA | TT5 | Huang lab |
| *Blautia hansenii* DSM 20583 | TT6 | Michael Fischbach's lab, Stanford |
| *[Clostridium] scindens* ATCC 35704 | TT7 | Sonnenburg lab |
| *[Clostridium] hylemonae* DSM 15053 | TT9 | Michael Fischbach's lab, Stanford |
| *Enterococcus faecalis* TX1322 | TT10 | Huang lab |
| *Hungatella hathewayi* WAL-18680 | TT11 | Huang lab |
| *Bacteroides fragilis* NCTC 9343 | TT12 | Sonnenburg lab |
| *Flavonifractor plautii* 1_3_50AFAA | TT13 | Huang lab |
| *Bacteroides uniformis* ATCC 8492 | TT14 | Michael Fischbach's lab, Stanford |
| *Parabacteroides distasonis* ATCC 8503 | TT15 | Michael Fischbach's lab, Stanford |
| *Salmonella enterica* serovar Typhimurium SL1344 | NA | Denise Monack's lab, Stanford |
| *Salmonella enterica* serovar Typhimurium 14028s pFVP25-mCherry | NA | Denise Monack's lab, Stanford |

**Table S3: Reference strains used in this study.**
